# Hypoxia triggers collective aerotactic migration in *Dictyostelium discoideum*

**DOI:** 10.1101/2020.08.17.246082

**Authors:** O. Cochet-Escartin, M. Demircigil, S. Hirose, B. Allais, P. Gonzalo, I. Mikaelian, K. Funamoto, C. Anjard, V. Calvez, J.-P. Rieu

**Affiliations:** Institut Lumière Matière, UMR5306, Université Lyon 1-CNRS, Université de Lyon, 69622 Villeurbanne, France; Institut Camille Jordan, UMR5208, Université Lyon 1-CNRS, Université de Lyon, 69622 Villeurbanne, France; Graduate School of Biomedical Engineering, Tohoku University, 6-6-12 Aramaki-aza Aoba, Aoba-ku, Sendai, Miyagi 980-8579, Japan; Institute of Fluid Science, Tohoku University, 2-1-1 Katahira, Aoba-ku, Sendai, Miyagi 980-8577, Japan; Centre Léon Bérard, Centre de recherche en cancérologie de Lyon, INSERM 1052, CNRS 5286, Université Lyon 1, Université de Lyon, 69373, Lyon, France; Laboratoire de Biochimie et Pharmacologie, Faculté de médecine de Saint-Etienne, CHU de Saint-Etienne, 42000, Saint-Etienne, France; Graduate School of Engineering, Tohoku University, 6-6 Aramaki-aza Aoba, Aoba-ku, Sendai, Miyagi 980-8579, Japan

## Abstract

It is well known that eukaryotic cells can sense oxygen (O_2_) and adapt their metabolism accordingly. It is less known that they can also move towards regions of higher oxygen level (aerotaxis). Using a self-generated hypoxic assay, we show that the social amoeba *Dictyostelium discoideum* displays a spectacular aerotactic behavior. When a cell colony is covered by a coverglass, cells quickly consume the available O_2_ and the ones close to the periphery move directionally outward forming a dense ring keeping a constant speed and density. To confirm that O_2_ is the main molecular player in this seemingly collective process, we combined two technological developments, porphyrin based O_2_ sensing films and microfluidic O_2_ gradient generators. We showed that *Dictyostelium* cells exhibit aerotactic and aerokinetic (increased speed at low O_2_) response in an extremely low range of O_2_ concentration (0-1.5%) indicative of a very efficient detection mechanism. The various cell behaviors under self-generated or imposed O_2_ gradients were modeled with a very satisfactory quantitative agreement using an *in silico* cellular Potts model built on experimental observations. This computational model was complemented with a parsimonious ‘Go or Grow’ partial differential equation (PDE) model. In both models, we found that the collective migration of a dense ring can be explained by the interplay between cell division and the modulation of aerotaxis, without the need for cell-cell communication.

## Introduction

Oxygen is the main electron acceptor for aerobic organism to allow efficient ATP synthesis. This high-energy metabolic pathway has contributed to the emergence and diversification of multicellular organism (Chen et al., 2015). While high O_2_ availability in the environment seems a given, its rapid local consumption can generate spatial and temporal gradients in many places, including within multicellular organism. Oxygen level and gradients are increasingly recognized as a central parameter in various physiopathological processes (Tonon et al., 2019), cancer and development. The well-known HIF (hypoxia inducible factor) pathway allows cells to regulate their behavior when exposed to hypoxia. At low O_2_ levels, cells then accumulate HIFα leading to the expression of genes that support cell functions appropriate to hypoxia (Pugh & Ratcliffe, 2017).

Another strategy used by organisms facing severe oxygen conditions is to move away from hypoxic regions, a mechanism called aerotaxis first described in bacteria (Engelmann, 1881; Winn, Bourne, & Mitchell, 2013). Aerotaxis will occur at the interface between environments with different oxygen content, such as soil/air, water/air or even within eukaryotic multicellular organisms between different tissues (Lyons, Reinhard, & Planavsky, 2014). In such organisms, oxygen was proposed to be a morphogen as in placentation (Genbacev, Zhou, Ludlow, & Fisher, 1997) or a chemoattractant during sarcoma cell invasion (Lewis et al., 2016) but such studies are relatively scarce due to the difficultly to measure oxygen gradients *in vivo*.

Recently, it was demonstrated that after covering an epithelial cell colony by a coverglass non permeable to O_2_, peripheral cells exhibit a strong outward directional migration to escape hypoxia from the center of the colony (Deygas et al., 2018). This is a striking example of collective response to a self-generated oxygen gradient. Self-generated chemoattractant gradients have been reported to trigger the dispersion of melanoma cells (Muinonen-Martin et al., 2014; Stuelten, 2017) or *Dictyostelium* cells (Tweedy, Susanto, & Insall, 2016) or the migration of the zebrafish lateral line primordium (Donà et al., 2013; Venkiteswaran et al., 2013). The mechanism is simple and very robust: the cell colony acts as a sink for the chemoattractant, removes it by degradation or uptake creating a gradient that, in turn, attracts the cells as long as the chemoattractant is present in the environment.

*Dictyostelium discoideum* (*Dd*) is an excellent model system to study the fairly virgin field of aerotaxis and of self-generated aerotaxis. *Dd* is an obligatory aerobic organism that requires at least 5% O_2_ to grow at optimal exponential rate (Cotler & Raper, 1968; Sandonà, Gastaldello, Rizzuto, & Bisson, 1995) while slower growth can occur at 2% O_2_. However, its ecological niche in the soil and around large amount of bacteria can result in reduced O_2_ availability. During its multicellular motile stage, high oxygen level is one of the signal used to trigger culmination of the migrating slug (Xu, Wang, Green, & West, 2012). For many years, *Dd* has been a classical organism to study chemotaxis and has emulated the development of many models of emergent and collective behavior since the seminal work of Keller and Segel (Hillen & Painter, 2009; Keller & Segel, 1970). An integrated approach combining biological methods (mutants), technological progress, and mathematical modeling is very valuable to tackle the issue of aerotaxis.

In this article, we study the influence of O_2_ self-generated gradients on *Dd* cells. Using first a simple confinement assay, then microfluidic tools, original oxygen sensors and theoretical approaches, we show that oxygen self-generated gradients can direct a seemingly collective migration of a cell colony. Our results confirm the remarkable robustness and long-lasting effect of self-generated gradients in collective migration. This case where oxygen is the key driver also suggests that self-generated gradients are widespread and a possible important feature in multicellular morphogenesis.

## Results

### Confinement triggers formation and propagation of a self-sustained cell ring

To trigger hypoxia on a colony of *Dd* cells, we used a vertical confinement strategy (Deygas et al., 2018). A spot of cells with a radius of about 1mm was deposited and covered by a larger glass coverslip with a radius of 9mm. We measured the vertical confinement through confocal microscopy and found the height between the bottom of the plate and the coverslip to be 50μm (Fig. S1).

Using spots containing around 2,000 cells (initial density around 10^3^ cells/mm^2^), the formation of a dense ring of cells moving outwards was observed as quickly as 30min after initiation of the confinement (Fig. 1A and movie M1). This formation time however depended non-linearly on initial cell density (the denser, the faster, Fig. S2). Once triggered, this collective migration was self-maintained for tens of hours, even days and the ring could, at these points, span centimeters (Fig. 1B).

**Fig1.**
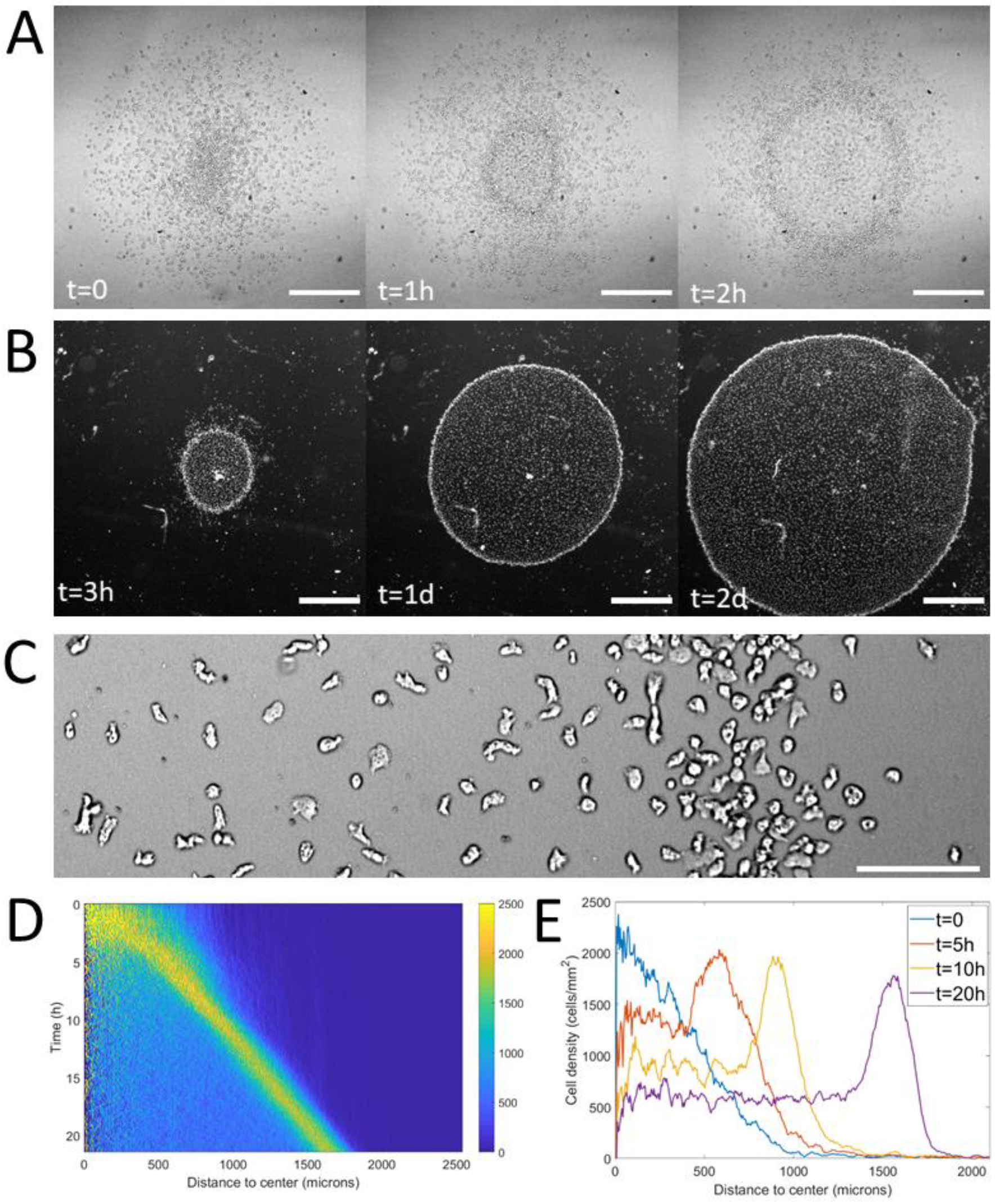
Formation and dynamics of a dense ring of cells after vertical confinement. (A) snapshots of early formation, scale bars: 500μm. (B) snapshots at longer times imaged under a binocular, scale bars: 1mm. (C) close up on a ring already formed moving rightward and showing different cellular shapes in the ring and behind it, scale bar: 100μm. (D) kymograph of cell density over 20h showing the formation and migration of the highly dense ring. (E) Cell density profiles in the radial direction at selected time points.

Notably, as the ring expanded outwards, it left a trail of cells behind. This led to the formation of a central zone populated by cells which didn’t contribute to the migration of the ring (Fig. 1B) but were still alive and moving albeit a clear elongated phenotype resembling pre-aggregative *Dd* cells (Fig. 1C and movie M2). In comparison, cells in the ring or outside the colony were rounder, as usual vegetative cells (Delanoë-Ayari et al., 2008).

To study the properties of the ring, we computed density profiles using radial coordinates from the center of the colony to study cell density as a function of time and distance to the center (Fig. 1D-E). We found that after a transitory period corresponding to the ring passing through the initial spot, the density in the ring, its width and its speed all remained constant over long time scales (Fig. S3). The speed and density of the ring were found to be 1.2 ± 0.3 μm/min (mean±std, N=9) and 1.9 10^3^ ± 0.3 10^3^ cells/mm^2^ (mean±std, N=4, *i.e.*, 4-fold higher than behind it, Fig. 1E) respectively. The density of cells left behind the ring was also found to remain constant after a transient regime (Fig. 1D). As the diameter of the ring increased over time, the absence of changes in morphology implies an increase of the number of cells and thus an important role of cell division.

Overall, this self-sustained ring propagation is very robust and a long lasting collective phenotype that can easily be triggered experimentally.

### Cell dynamics in different regions

Following the reported shape differences, we questioned how cells behaved dynamically in different regions. To do so, we performed higher resolution, higher frame rate experiments to allow cell tracking over times on the order of tens of minutes. Both the cell diffusion constant and instantaneous cell speeds were fairly constant throughout the entire colony (Fig. S4). Cell diffusion was 28.2 ± 1.4 μm^2^/min (N=3), comparable to our measurement of activity at very low oxygen level in the microfluidic device (see below). To test the influence of motion bias, we projected cell displacements on the radial direction and computed mean speeds in this direction as a function of distance to the center. Random motion, either persistent or not, would lead to a null mean radial displacement whereas biased migration would be revealed by positive (outward motion) or negative (inward motion) values. Here, we found that significantly non-zero biases were observed only in a region spanning the entire ring and a few tens of microns behind and in front of it with the strongest positive biases found in the ring (Fig. S5).

Overall, our results show that the different regions defined by the ring and its dynamics can be characterized in terms of cell behavior: (i) behind the ring in the hypoxic region: elongated shape, normal speeds, and low bias; (ii) in the ring: round shape, normal speeds and high bias.

### Response of *Dd* cells to controlled oxygen gradients

The spot assay is experimentally simple but is not ideally suited to decipher the response of *Dd* cells to hypoxia since local concentrations and gradients of oxygen are coupled to cell dynamics and thus very difficult to manipulate. We hence designed a new double-layer PDMS microfluidic device allowing to quickly generate oxygen gradients in a continuous, controlled manner (Fig. 2A). Briefly, cells were seeded homogenously within a media channel positioned 500μm below two gas channels continuously perfused with pure nitrogen on one side and air on the other. As PDMS is permeable to these gases, the gas flows imposed hypoxic conditions on one side of the media channel while the other was kept at atmospheric oxygen concentration. Of note, the distance between the two gas channels, thereafter called the gap, varied from 0.5mm to 2mm in order to modify the steepness of the gradients in the median region of the media channels (Fig. 2A and supplementary information).

**Fig. 2.**
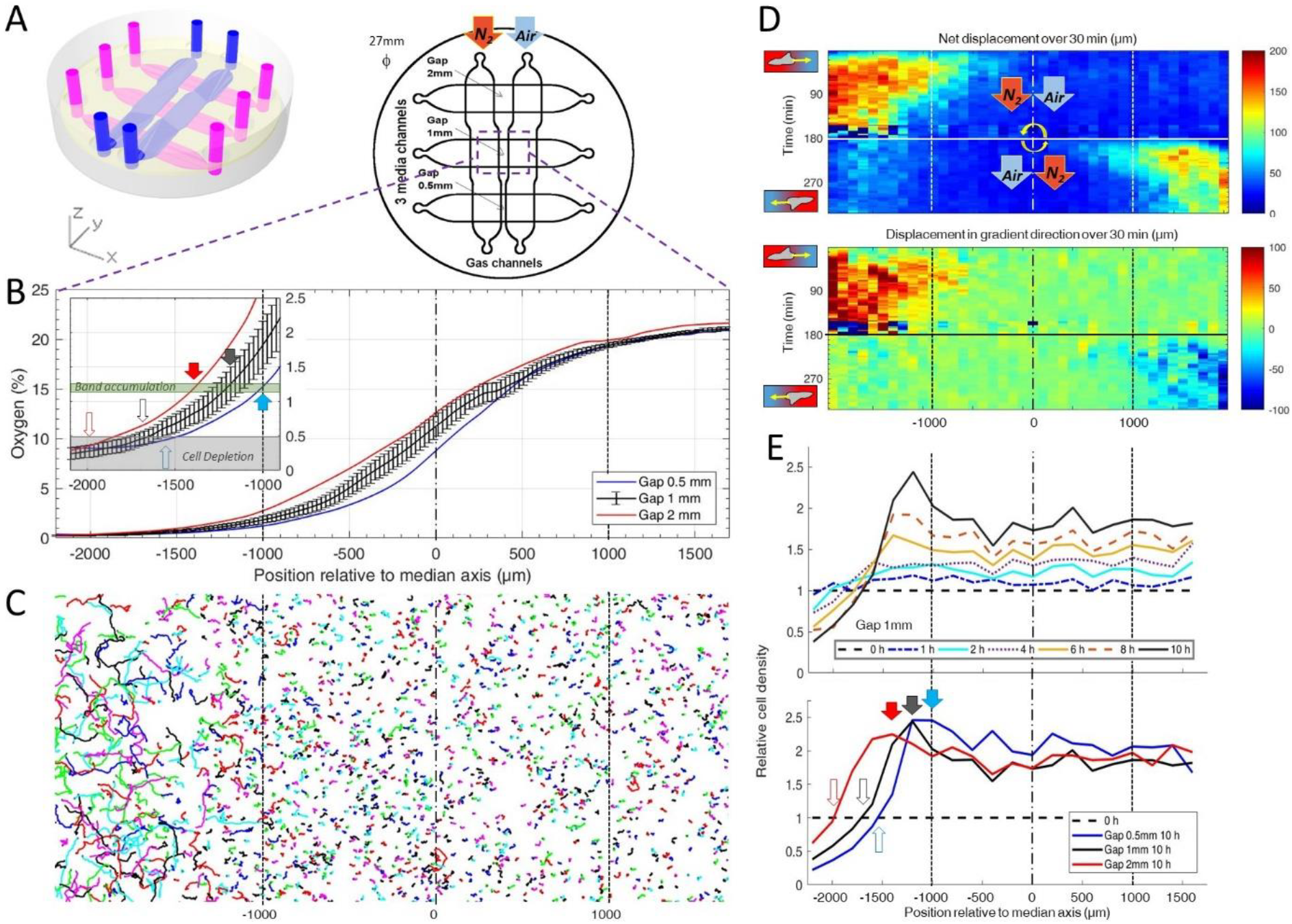
Dictyostelium single cells are attracted by an external O_2_ gradient when O_2_ level drops below 2%. (A) Schemes of the new double-layer PDMS microfluidic device allowing the control of the O_2_ gradient by the separation distance (gap) between two gas channels located 0.5 mm above the three media channels and filled with pure nitrogen, and air (21% O_2_). (B) Measured O_2_ concentration profiles 30 min after N2-Air injection to the left and right channels respectively (0-21% gradient) as a function of the position along the media channel for the three gaps. Error bars (see supplementary information) are reported only for gap 1 mm for clarity. The inset shows the 0%-2.5% region under the nitrogen gas channel (arrows, see E). (C) Trajectories lasting 1 h between 3h and 4h after establishment of a 0-21% gradient. Cells are fast and directed toward the air side in the region beyond the −1000 μm limit (O_2_<2%). (D) Cell net displacement over 30 min (end to end distance, top kymograph) and 30-min displacement projected along gradient direction (bottom kymograph). Cells are fast and directed toward O_2_, where O_2_<2%, within 15 min after 0-21% gradient establishment at t=0. At t=180 min, the gradient is reversed to 21-0% by permuting gas entries. Cells within 15 min again respond in the 0-2% region. (E) Relative cell density histogram (normalized to t=0 cell density) as a function of the position along media channel. Top panel: long term cell depletion for positions beyond −1600 μm (O_2_<0.5%, see inset of B) and resulting accumulation at about −1200 μm for gap 1 mm channel. The overall relative cell density increase is due to cell divisions. Bottom panel: cell depletion and accumulation at 10 h for the three gaps. The empty and filled arrows pointing the limit of the depletion region, and max cell accumulation respectively are reported in the inset of B).

To make sure that the gas flows were sufficient to maintain a constant O_2_ distribution against leakages and against small variation in the fabrication process, we also developed O_2_ sensing films to be able to experimentally measure O_2_ profiles both in the microfluidic devices and in the spot assay. These films consisted of porphyrin based O_2_ sensors embedded in a layer of PDMS. As O_2_ gets depleted, the luminescence quenching of the porphyrin complex is reduced leading to an increase in fluorescence intensity (Ungerböck, Charwat, Ertl, & Mayr, 2013). Quantitative oxygen measurements were then extracted from this fluorescent signal using a Stern-Volmer equation (See supplementary information and Figs. S6-S9 for details).

First, we observed the formation of a stable O_2_ gradient in the devices closely resembling numerical predictions with or without cells (Fig. 2B and Fig. S10-12).

We then turned our attention to the reaction of the cells to this external gradient. We first noticed that depending on local O_2_ concentrations, cell motility was remarkably different. Using cell tracking, we found that cell trajectories seemed much longer and more biased in hypoxic regions (Fig. 2C). These aerokinetic (large increase in cell activity) and aerotactic responses were confirmed by quantifying the mean absolute distance travelled by cells (Fig. 2D top), or the mean distance projected along the gradient direction (Fig. 2D bottom) in a given time as a function of position in the device (Fig. 2D).

The second important observation stemming from the microfluidic experiments is an accumulation of cells at some midpoint within the cell channel (Fig. 2E). Naively, one could have expected cells to follow the O_2_ gradient over its entire span leading to an accumulation of all cells on the O_2_ rich side of the channel. This did not happen and, instead, cells seemed to stop responding to the gradient at a certain point. Similarly, we observed a strong positive bias in hypoxic regions but the bias quickly fell to 0 as cells moved to oxygen levels higher than about 2% (Fig. 2D), confirming that the observed cell accumulation was a result of differential migration and not, for example, differential cell division. In addition, if we inverted the gas channels halfway through the experiment, we observed that the cells responded in around 15min (which is also the time needed to re-establish the gradient, see Fig. S11) and showed the same behavior, albeit in reverse positions. We measured the bias for the different gaps and for the situation of reversed gradient and obtained a value of 1.1 ± 0.4 μm/min (N=6).

Of note, the position at which cells accumulated and stopped responding to the gradient was still in the region were the gradient was constantly increasing. This led to the hypothesis that, in addition to gradient strength, O_2_ levels also play an important role in setting the strength of aerotaxis displayed by *Dd* cells.

Furthermore, when we compared experiments performed with different gaps, we found that the position of cell accumulation varied from one channel to another (Fig. 2E). However, our O_2_ sensors indicates that the accumulation occurred at a similar O_2_ concentration of about 1% in all three channels (inset of Fig. 2B) thus strongly hinting that the parameter controlling the aerotactic response was O_2_ levels.

Overall, these experiments in controlled environments demonstrated two main features of the response of *Dd* cells to hypoxia: a strong aerokinetic response and a positive aerotactic response, both modulated by local O_2_ levels regardless of the local gradient. These results reveal a subtle cross talk between O_2_ concentrations and gradients in defining cell properties and it would be very informative, in the future, to study in details the reaction of *Dd* cells to various, well defined hypoxic environments where O_2_ concentrations and gradients can be independently varied.

### Coupled dynamics between oxygen profiles and collective motion

Thanks to these results, we turned our attention back to the collective migration of a ring of cells and asked whether similar aerotactic behaviors were observed under self-generated gradients. To do so, we performed spot experiments on the O_2_ sensing films described above which allowed us to image, in parallel, cell behavior and O_2_ distribution (Figs. 3A and S8, Movie M3).

**Fig3.**
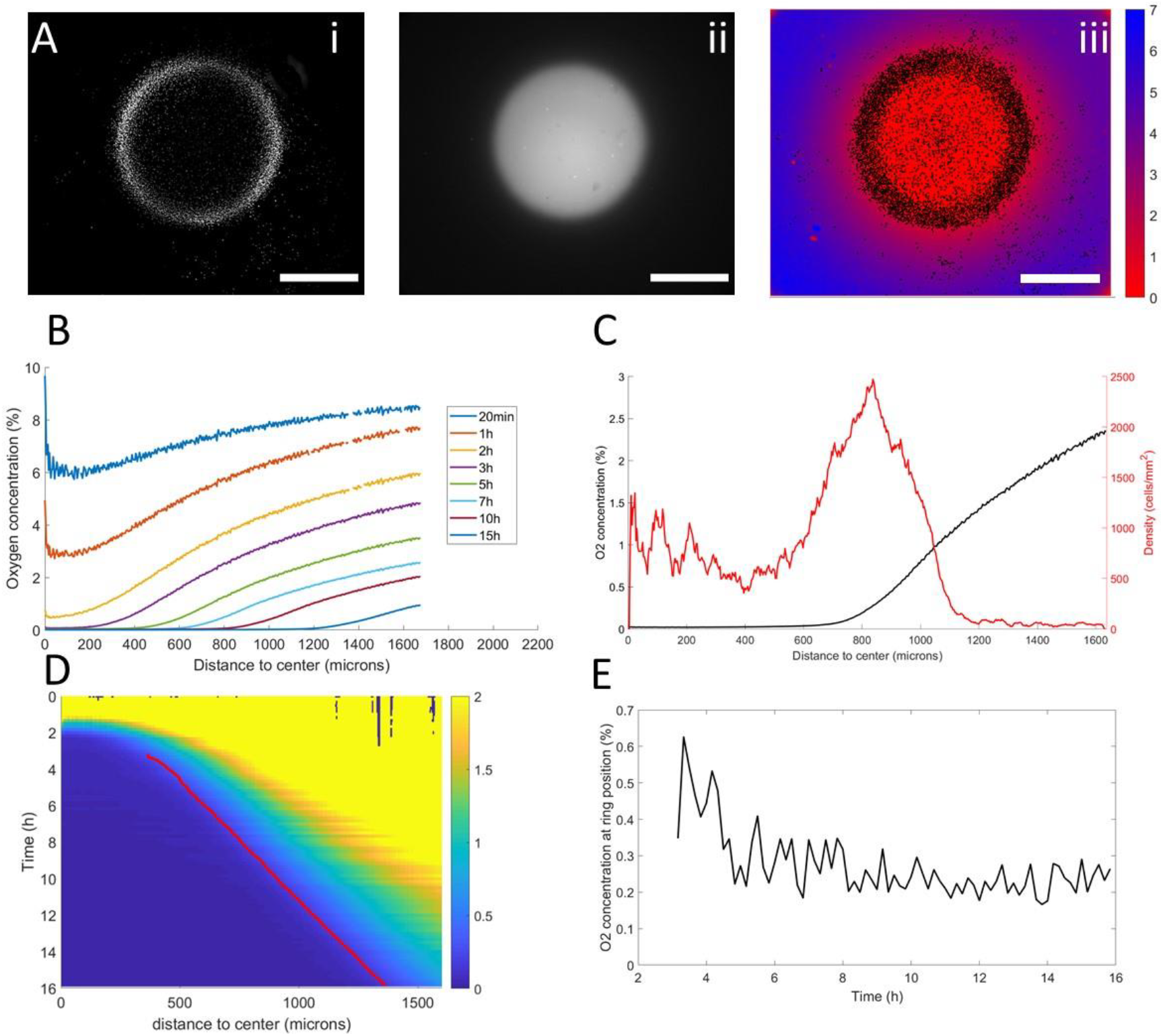
Interplay between ring dynamics and O_2_ profiles. (A) i: treated image showing cell distribution at t=10h, ii: raw fluorescent signal indicative of strong O_2_ depletion, iii: reconstructed image showing the center of mass of all detected cells and quantitative O_2_ profiles (colorbar, in % of O_2_), scale bars: 1mm. (B) O_2_ profiles averaged over all angles and shown at different times. (C) radial profile of cell density and O_2_ concentration at t=10h showing the position of the ring relative to the O_2_ profile. (D) kymograph of O_2_ concentration (colorbar in %) with the position of the ring represented as a red line. The colormap is limited to the 0-2% range for readability but earlier time points show concentrations higher than the 2% limit. (E) O_2_ concentration as measured at the position of the ring as a function of time showing that the ring is indeed following a constant concentration after a transitory period.

In a first phase, preceding the formation of the ring, cell motion was limited and the structure of the colony remained mostly unchanged. As O_2_ was consumed by cells, depletion started in the center and sharp gradients appeared at the edges of the colony (Fig. 3B-C).

Then, the ring formed and started moving outwards, O_2_ depletion continued and the region of high O_2_ gradients naturally started moving outwards (Fig. 3B). At this point, coupled dynamics between the cells and the O_2_ distribution appeared and we observed that the position of the ring closely followed the dynamics of the O_2_ field (Fig. 3D), *i.e.* a constant concentration of oxygen of less than 1% (Fig. 3C).

In the process, three distinct regions were created. Behind the ring, O_2_ was completely depleted and thus no gradient was visible. In front of the ring, the O_2_ concentration remained high with high gradients. Finally, in the ring region, O_2_ was low (<1%) and the gradients were strong. Based on our results in externally imposed gradients, we would thus expect cells to present a positive aerotactic bias mostly in the ring region which is indeed what we observed (Fig S5).

### Minimal cellular Potts Model

Based on these experimental results, we then asked whether this subtle response of *Dd* cells to complex oxygen environments was sufficient to explain the emergence of a highly stable, self-maintained collective phenomenon. To do so, we developed cellular Potts models based on experimental observations and tested whether they could reproduce the observed cell dynamics. Briefly, the ingredient underlying the model are as follows (details can be found in the supplementary information). First, all cells consume the oxygen that is locally available at a known rate (Torija et al., 2006). Cell activity increases at low O_2_. Cells respond positively to O_2_ gradients with a modulation of the strength of this aerotaxis based on local O_2_ concentrations, as observed in our microfluidic experiments. Finally, all cells can divide. Of note, all parameters were scaled so that both time and length scales in the Potts models are linked to experimental times and lengths (see Supplementary Information).

Although this model is based on experimental evidence, some of its parameters are not directly related to easily measurable biological properties. Therefore, we decided to fit our parameters to reproduce as faithfully as possible the results of our microfluidic experiments. Through a trial and error procedure, we managed to reproduce these results qualitatively and quantitatively (Movie M4) in terms of collective behavior, cell accumulation, and individual cell behavior (Fig. S13).

We then applied this model and added O_2_ consumption by cells, with initial conditions mimicking our spot assay and other ingredients mimicking the vertical confinement (see Supplementary Methods). We observed the rapid formation and migration of a ring (Fig. 4A-B, Movie M5). This ring was remarkably similar to that observed in experiments. In particular, we found its speed to be constant after an initial transitory period (Fig. 4C, Fig. S14). This speed was also comparable to experimental ones. Similarly, the morphology of these simulated rings was constant over time with a fixed cell density and width (Fig. S14). Finally, cell behavior was qualitatively well reproduced by this model (Fig. S15) with the exception of a remaining dense core of cells in the center which disappeared if cell consumption (or initial cell density) was reduced (see Fig 5A).

**Fig. 4.**
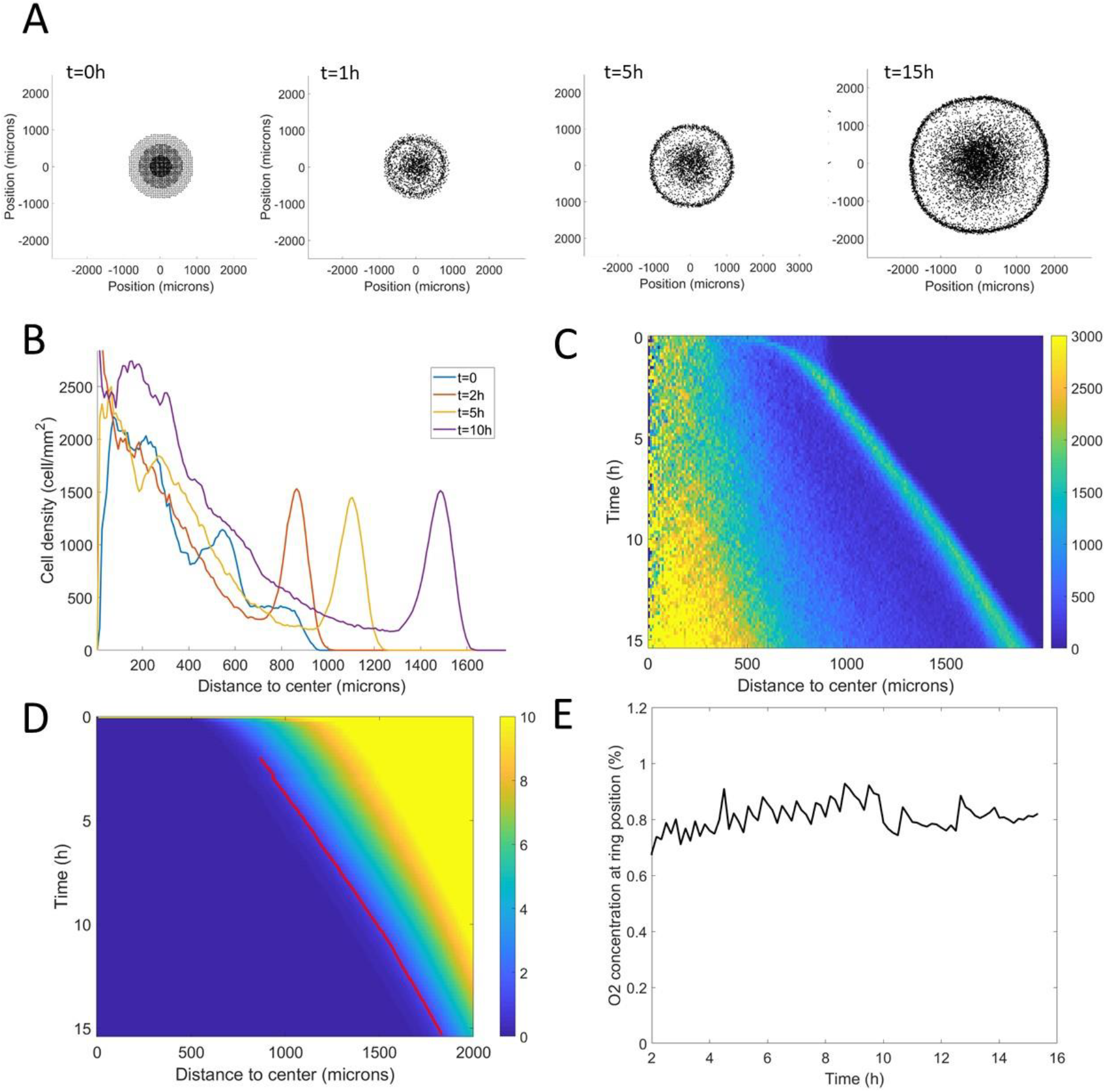
Minimal Potts model of ring formation and migration. (A) Snapshots of a simulated colony of cells showing the formation of highly dense ring of cells. (B) Cell density profiles averaged over all angles for four different times. (C) Corresponding kymograph of cell density (colorbar in cells/mm^2^) as a function of time and distance to the center. Quantification in terms of microns and hours is described in the supplementary information. (D) Kymograph of O_2_ concentration (colorbar in %) with the position of the ring represented as a red line. The colormap is limited to the 0-10% range for readability but earlier time points show concentrations higher than the 10% limit. (E) O_2_ concentration at the ring position as a function of time showing that, here too, the ring follows a constant O_2_ concentration.

**Fig. 5.**
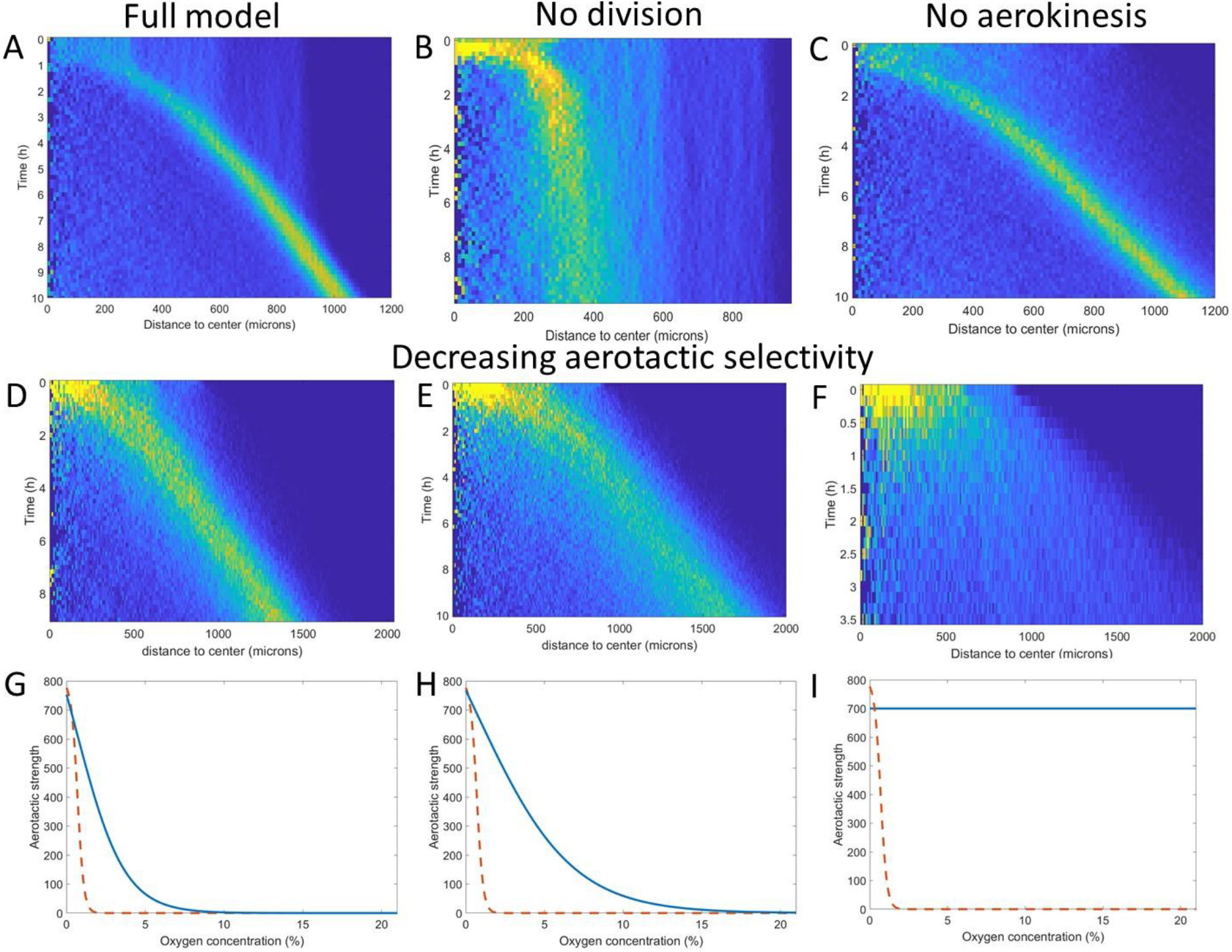
Key ingredients of the Potts model by density kymograph (DK) evaluation. (A) DK for the full model with reduced oxygen consumption as a basis for comparison. (B) DK in the absence of cell division, note the difference in length scale showing a clear limitation of motion in that case. (C) DK in the absence of aerokinesis (cell activity is not modulated by local oxygen concentrations). (D) DK with a modulation of aerotactic strength as shown in (G), note the wider ring. (E) DK with a modulation of aerotactic strength as shown in (H). (F) DK with a modulation of aerotactic strength as shown in (I), no ring appears and cells quickly migrate outwards as shown by the difference in time scales. (G-I) Three different aerotactic modulations, in blue, compared to the one used in the full model, shown in (A), drawn here as a red dashed line.

In terms of coupled dynamics between cell density and O_2_ profiles, we found here too that the driving force behind this collective phenomenon was the fact that the ring followed a constant O_2_ concentration (Fig. 4D-E).

We then asked what were the key ingredients in the model to trigger this phenomenon, a question we explored by tuning our original Potts model. We started by dividing cell consumption of oxygen by a factor of 3 (Fig. 5A) and found that it didn’t significantly change the ring speed but could change the aspect of cell density in the central region. We then turned our attention to other key elements in the model.

If we turned off cell division in our models, the formation of the ring was mostly unchanged but after a short time, the ring started slowing down and even stopped as cell density was no longer sufficient to reach highly hypoxic conditions (comparing Fig5 A and B). Second, we asked whether the observed and modelled aerokinesis was necessary to reproduce the collective migration. We found that it wasn’t as models ran at different effective temperatures applied to all cells regardless of local O_2_ concentrations, all showed qualitatively similar behavior (see for example Fig. 5C). Of note though, lower effective temperatures led to less dense rings as fewer cells were able to start in the ring (Fig. S16). Finally, we found that modulation of aerotaxis by local O_2_ concentrations was essential. Indeed, as we increased the range of O_2_ concentration at which aerotaxis is at play (Fig 5G-H), we found that forming rings became wider and less dense (Fig. 5 D-E) to the point where no actual ring could be distinguished if aerotaxis was kept constant for all cells (Fig. 5I).

These numerical simulations based on cellular Potts models provide a good intuition of the phenomenon and reveal that cell division and aerotactic modulation are the two key ingredients to reproduce the ring of cells. They fall short however of giving an in-depth quantitative description because they rely on many parameters and are not amenable to theoretical analysis *per se*.

### Single-threshold ‘Go or Grow’ model

For the above mentioned reasons, we developed a simple, yet original, model based on the two key ingredients while keeping it analytically solvable. Cell density is treated as a continuous field, O_2_ consumption and diffusion are included. Cells have two distinct behaviors, depending on the O_2_ concentration. Below a certain threshold *C*_0_ cells move preferentially upward the oxygen gradient, with constant advection speed *a*_0_, but they cannot divide. Above the same threshold they divide and move randomly without directional bias. We refer to these behaviors as respectively *go* or *grow*. We thereby revisit the ‘Go or Grow’ model for glioma cells (Hatzikirou, Basanta, Simon, Schaller, & Deutsch, 2012) with the transition between division and directional motion being mediated by oxygen levels rather than cell density. This model is meant to 1-demonstrate that these ingredients suffice to trigger a collective motion and 2-determine the relative contributions of division and aerotaxis on the speed of the ring.

For the sake of simplicity and its relevance for the study of planar front propagation, we restrict ourselves to the one-dimensional case, ignoring curvature effects. Our ‘Go or Grow’ model is a piecewise constant reaction-advection-diffusion equation:

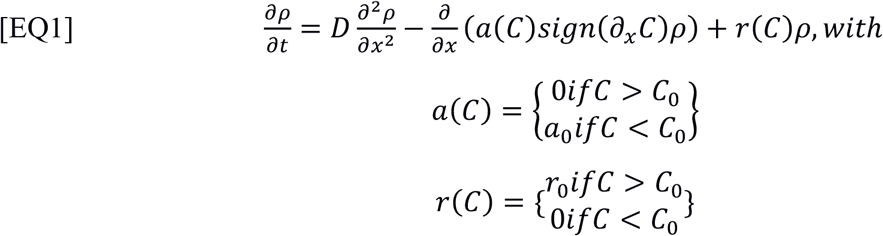

*a*(*C*)*sign*(∂_*x*_*C*) corresponds to the aerotactic advection speed and *r*(*C*) to the cell division rate. Oxygen is subject to a simple diffusion-consumption equation, with *b* the constant consumption rate of oxygen per cell:

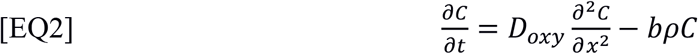

The coupling goes through the location of the oxygen threshold *C*_0_.

We investigated the existence of traveling wave profiles at some unknown speed σ > 0. The equation for the one-dimensional profile, being stationary in the moving frame *z* = *x* − σ*t*, is:

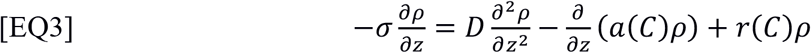

Interestingly enough, it admits explicit solutions, whose speed does not involve the dynamics of oxygen consumption. This yields the following formula for the wave speed (Fig. 6A):

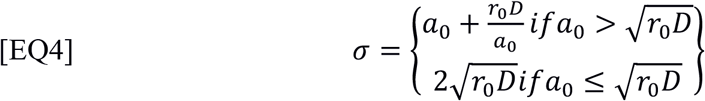

**Fig. 6.**
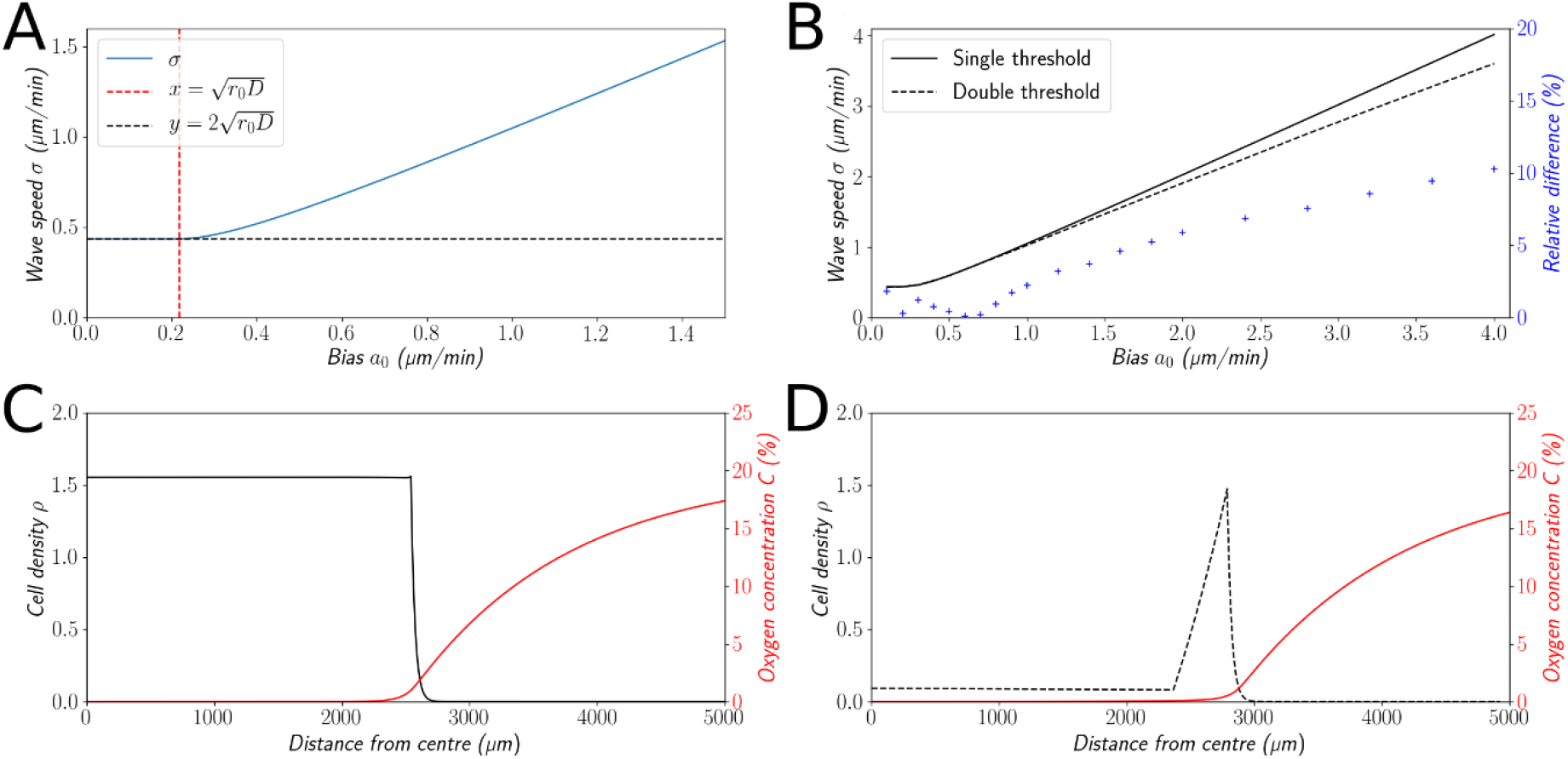
Simple ‘Go or Grow’ model. (A): Illustration of wave speed given by EQ4. (B): Comparison of wave speeds for the single-threshold and the double-threshold models (solid and dotted lines respectively) and relative difference between the two models (crosses). (C): Traveling wave profile of cell density and O_2_ concentration for the single-threshold model (EQ1 & EQ2). (D): Traveling wave profile of cell density and O_2_ concentration for the double-threshold model. In both cases, thresholds coincide with points of sharp transitions in the profiles.

To the best of our knowledge, this analytical formula is new and captures basic features of a wave under a single self-generated gradient.

### Double-threshold ‘Go or Grow’ model and variations

The cell density profile (Fig. 6C) of the ‘Go or Grow’ model EQ1 is clearly not in agreement with the observation of a dense ring of cells followed by an expanding zone at lower density (Fig. 1). Yet through elementary structural variations of the model, we obtained cell density profiles more similar to experimental ones, without significant alterations to the quantitative and qualitative conclusions.

First, we made the hypothesis of a second oxygen threshold *C*_0_′ < *C*_0_, below which cells are not sensitive to gradients any longer. Notably, this model exhibits traveling wave profiles with a dense ring of cells, leaving behind a spatially uniform low cell density (Fig. 6D) much more similar to experimental observations. An exhaustive analytical investigation of this model seems out of reach, but we analyzed it numerically. The speed of the numerical wave remained close to the value given by formula EQ4 (at most 10% of relative difference, Fig. 6B). Intuitively, the main contribution to the collective speed is the strong bias inside the ring at intermediate levels of O_2_, whereas cells at levels below the second threshold *C*_0_′, where the dynamics of both models diverge, do not contribute much to the collective speed.

Second, we investigated more complex responses of cells to oxygen levels (Fig. S17). We assumed a continuous dependency of the bias to the O_2_ concentration, rather than a discontinuous switch as in EQ1. In parallel, we assumed a continuous dependency of the division rate. The conclusions remained qualitatively and quantitatively the same (Fig. S17).

Therefore, properties of the ‘Go or Grow’ model, which we can prove analytically, like formula EQ4 and features discussed below, appear to be robust to substantial variations.

### Comparison with the Fisher-KPP reaction-diffusion equation

The Fisher-KPP equation is a standard model for the linear expansion of populations (Fisher, 1937; Kolmogorov, Petrovskii, & Piskunov, 1937). It describes front propagation under the combined effects of diffusion and growth. The wave speed of our ‘Go or Grow’ model coincides with Fisher’s speed, i.e. 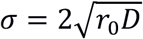, in the regime of small bias 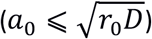. This is the signature of a *pulled wave*, meaning that the propagation is driven by the division and motion of cells at the edge of the front, with negligible contribution from the bulk. In contrast, when the bias is large 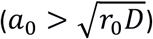 then the wave speed in [EQ4] is greater than Fisher’s speed. This is the signature of a *pushed wave*, meaning that there is a significant contribution from the bulk to the net propagation, see (Birzu, Hallatschek, & Korolev, 2018) for recent insights about the dichotomy between pulled and pushed waves.

In order to explore this dichotomy between pulled and pushed waves, we used the framework of neutral labeling (Roques, Garnier, Hamel, & Klein, 2012) in the context of PDE models. We colored fractions of the density profile to mimic labeling of cells with two colors. Then, we followed numerically the dynamics of these fractions, and quantified the mixing of the two colors. Our results are in perfect agreement with (Roques et al., 2012), extending their results beyond classical reaction-diffusion equations, including advection (see supplementary information section 3.2). In the case of large bias (Fig. 7A-C) the wave is pushed and the profile is a perfect mixture of blue and yellow cells at long time. Contrarily, the wave is pulled in the regime of small bias: only cells that were already initially in the front, here colored in blue (Fig. 7B-D), are conserved in the front, whilst yellow cells at the back cannot catch up with the front.

**Fig. 7.**
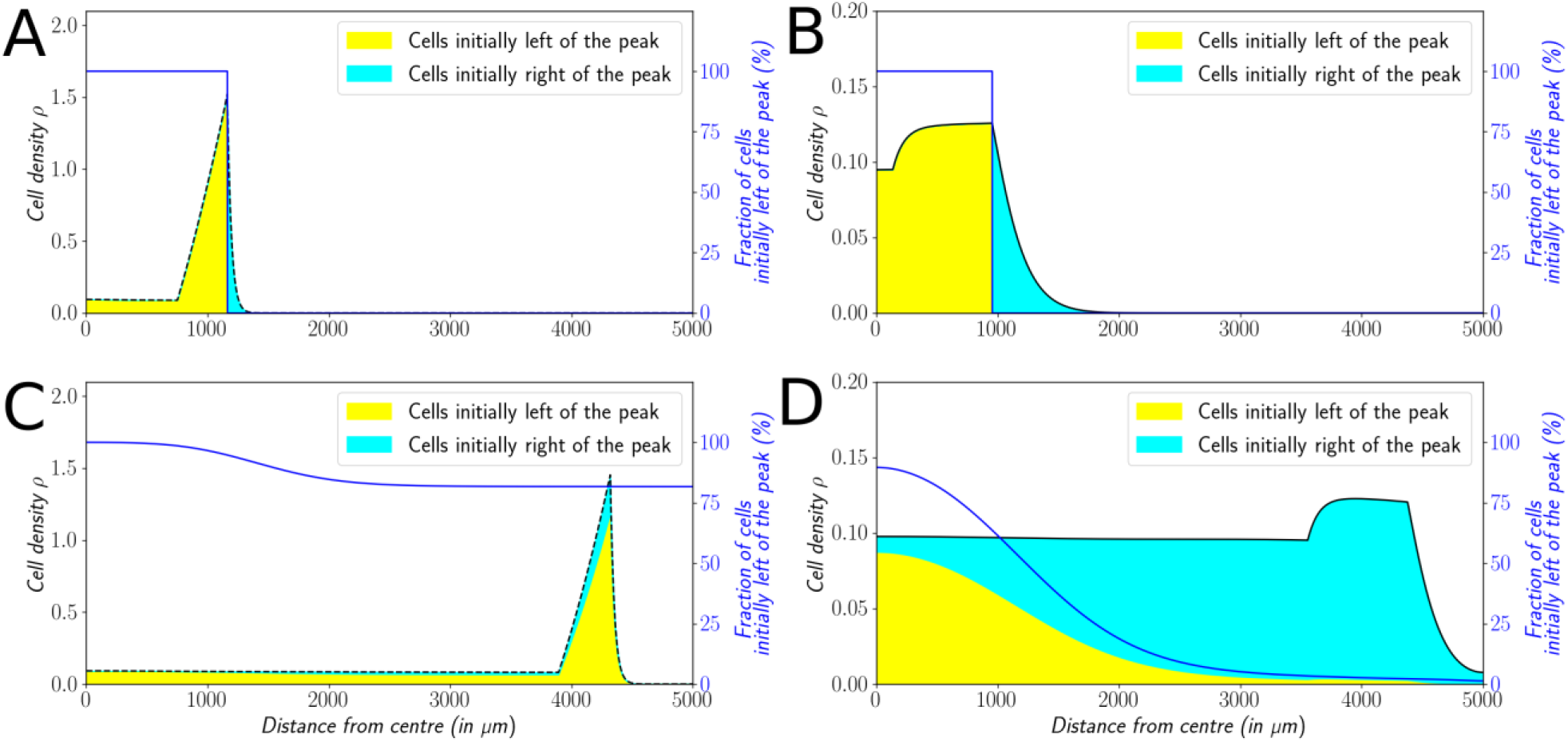
Classification of the expansion type. Cells initially on the left-hand side or right-hand side of the peak get labeled differently (A&B). The labeling is neutral and does not change the dynamics of the cells. We let evolve the two colored population for some time and observe the mixing of the colors (C&D). (A&C): With *a*_0_ = 1μ*m* ⋅ *s*^−1^, the wave is pushed wave and after some time the front undergoes a spatially uniform mixing. (B&D): With *a*_0_ = 0.1μ*m* ⋅ *s*^−1^, the wave is pulled and only the fraction initially in the front is conserved in the front.

In the absence of associated experimental data, we explored the cellular Potts model with such neutral labeling. The results were in agreement with the PDE simulations (Fig. S18) showing a clear, rapid mixing of the two cell populations under the propagation of a pushed wave.

### Relative contribution of growth and aerotactic bias to the collective motion in experiments

In [EQ4] we see that growth contributes to the wave speed either via the geometric mean of growth rate *r*_0_and diffusion coefficient *D*, in the case of small bias, or via a linear term supplementing the advection speed *a*_0_ for large bias. As a rough approximation with *a*_0_ = 1μ*m* ⋅ *min*^−1^, assuming a doubling time of 8h for *Dd* cells and *D* = 30μ*m*^2^ ⋅ *min*^−1^, we are clearly in the case of large bias 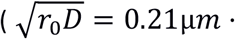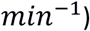 and EQ4 yields σ = 1.04μ*m* ⋅ *min*^−1^. The exact contribution of growth depends on the variations of the model (numerical explorations have shown contributions of growth of at most 20%, see Supplementary Information), yet even with this elementary reasoning we were able to estimate that growth contributes only up to a small fraction to the overall speed.

Overall, the invesvestigation of the ‘Go or Grow’ model and its variations show that the ring speed is mainly the outcome of the interplay between cell division and aerotaxis.

## Discussion

While aerotaxis is well established for bacteria, its role is often invoked in multicellular organisms to explain various processes in development or cancer progression, but very few *in vitro* studies were conducted to prove it is an efficient and operating mechanism or to understand the molecular mechanisms at play during aerotaxis. Deygas et al. showed that confined epithelial colonies may trigger a self-generated O_2_ gradient and an aerotactic indirect response through a secondary ROS self-generated gradient (Deygas et al., 2018). Gilkes et al. showed that hypoxia enhances MDA-MB231 breast cancer cell motility through an increased activity of HIFs (Gilkes et al., 2014). HIFs activate transcription of the Rho family member RHOA and Rho kinase 1 (ROCK1) genes, leading to cytoskeletal changes, focal adhesion formation and actomyosin contractions that underlie the invasive cancer cell phenotype. This study suggests a role for aerotaxis in tumor escape, but it only demonstrates aerokinesis as O_2_ gradients were not imposed to probe a directed migration toward O_2_. Using a microfluidic device, the same cancer cell line was submitted to various oxygen levels as well as oxygen gradients(Koens et al., 2020) but the observed aerotactic response was not clear.

By contrast, the experimental results presented here with *Dd* show a strong response to hypoxia. Within 15min, cells exhibit an aerokinetic and aerotactic response when exposed to externally imposed O_2_ gradients (Fig. 2). Self-generated O_2_ gradients are produced within 20 min (Fig. 3 and Fig. S2). But this cellular response is within the equilibration time of the oxygen distribution (Fig. S11). Hence we can consider the cellular response as almost instantaneous with *Dd*. The difference with previously studied cells is probably due to the extreme plasticity of the rapidly moving amoeboid cells (*Dd*) and their almost adhesion independent migration mechanism (Friedl, Borgmann, & Bröcker, 2001) while mesenchymal cancer cells move slower by coordinating cytoskeleton forces and focal adhesion (Palecek, Loftust, Ginsberg, Lauffenburger, & Horwitz, 1997).

The quick response of *Dd* in directed migration assays has been largely exploited to decipher the molecular mechanisms at play during chemotaxis (Nakajima, Ishihara, Imoto, & Sawai, 2014). The molecular mechanisms used for O_2_ sensing and its transduction into cellular response are for the moment unknown but we can expect that the O_2_ molecular sensors modulate cytoskeleton organization, particularly localized actin polymerization/depolymerization through some of the molecular components involved in classical chemotaxis toward folate or cAMP (Pan, Xu, Chen, & Jin, 2016; Van Haastert, Keizer-Gunnink, & Kortholt, 2017). However, new and unexpected mechanisms cannot be excluded.

The finding that migrating cells can influence the direction of their own migration by building chemoattractant gradients is not new but it was only recently reported for various systems (Stuelten, 2017): melanoma cells that break down lysophosphatidic acid (LPA) and generate a LPA gradient in the vicinity of the tumor (Muinonen-Martin et al., 2014), *Dd* colonies that generate folate gradients (Tweedy et al., 2016) or for the migration of the zebrafish lateral line primordium through a self-generated chemokine gradient (Donà et al., 2013; Venkiteswaran et al., 2013). The dispersal of melanoma cells is particularly instructive. The stroma surrounding the tumor acts as a source of LPA. The tumor cells act as a sink for LPA. As long as LPA is present in the environment a steady wave of migrating melanoma cells propagates away from the initial tumor over long distances and long time periods.

The self-generated LPA (melanoma) and folate (*Dd*) gradients were modeled with a simple numerical model that was able to predict the steady wave. In particular, it predicted an invasive front where cells are exposed to a steep chemoattractant gradient, followed by a ‘trailing end’ where the gradient is shallow and fewer cells migrate with poor directionality(Tweedy & Insall, 2020). It also predicted that the wave may have a less marked front, and/or a smaller speed, or even vanishes if the cell density was too low due to insufficient chemoattractant removal. All these features are surprisingly similar to our experimental measurements of cell density and O_2_ profiles (Fig. 1E, Fig. 3C). The atmospheric O_2_ that diffuses through the culture medium and eventually the plastic surfaces is the chemoattractant. The O_2_ consumption triggers hypoxia that in turn generate an aerotactic response toward O_2_ in a very narrow range of O_2_ concentrations (0-1.5%) (Fig. 3C). The exact value of the lower O_2_ threshold value will deserve future investigations. The exact nature of the cellular response at these extremely low O_2_ levels, and in a very shallow gradient, also has yet to be clarified.

Our different models unveil a set of basic assumptions which are sufficient for collective motion of cells without cell-cell interactions (attractive or otherwise), in contrast with (Saragosti et al., 2011). Cell growth is necessary to produce a long-standing wave without any damping effect. However, it may not be the main contribution in the wave speed, depending on the relative ratio between directional motion (the bias *a*_0_), and reaction-diffusion (the Fisher half-speed 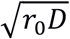). In the case where the former is greater than the latter, the wave is due to the combination of growth and directional motion and it is pushed. This result differs particularly from the Fisher equation with constant advection (meaning with uniform migration and division) where the wave speed is 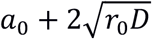 and the wave is pulled. In the experiments under study, we estimate directional motion to contribute the most to the cell speed, ruling out the possibility of seeing a pulled wave driven by cell division and diffusion at the edge of the front.

In conclusion, we demonstrate the remarkable stability of collective motion driven by self-generated gradients through depletion of an attractant. Through coupled dynamics, these gradients give rise to long lasting, communication-free migrations of entire colonies of cells which are important both from ecological and developmental points of view. In the case presented here where oxygen plays the role of the depleted attractant, this could prove to be a very general mechanism as oxygen is ubiquitous and always consumed by cells.

## Methods

### Cell preparation

Cells were grown in HL5 media at 22°C with shaking at 180 rpm for oxygenation (Sussman, 1987). Exponentially growing cells were harvested, counted to adjust cell density at typically 2000 cells/μL. For the spot assay, 1 μL of cell suspension was carefully deposited on a dry surface (6 wells plastic plates or PDMS for oxygen sensing) and incubated for 5 to 7 minutes in humid atmosphere. 2 ml HL5 was slowly added to avoid detaching cells and a wetted coverslip was then deposited on top of the micro-colony. The micro-colony was then observed under video-microscope and analysed with ImageJ and Matlab (see supplementary information).

### Microfluidic devices and O_2_ sensing films

Using soft lithography, we manufactured a double layer PDMS microfluidic device to control a O_2_ gradient along the cell-media channel. The gradient is established within 15 min. We also developed coverglass coated with a thin layer of PDMS with porphyrin embedded: the quenching by O_2_ of the porphyrin luminescence enables the direct measurement of the O_2_ concentration from a proper calibration of the fluorescence intensity (details are given in the supplementary information).

### Potts models

Potts model simulations were run using CompuCell3D (Swat et al., 2012) with a mix of prebuilt modules and home-made Python steppables in particular to implement the modulation of aerotactic strength by local oxygen levels. Most parameters were fitted to experimental measurements and both time and length scales were also adapted to achieve quantitative simulations. Full details of the model can be found in Supplementary Methods and full codes are available upon request to the corresponding authors.

### Go or grow model and simulations

Both diffusion equations EQ1&EQ2 were discretized through a time-backward space-centered difference scheme with an upwind discretization for the advection operator:

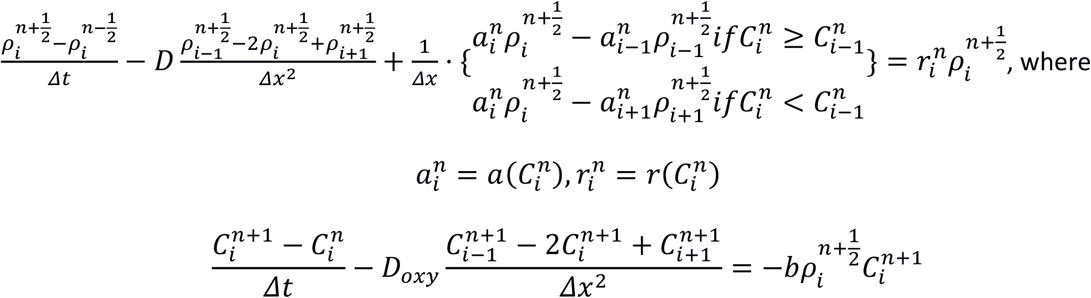

The schemes were coded in Python language. All simulations of EQ1&EQ2 shown in this article were carried out with a mesh size Δ*t* = 1*min* and Δ*x* = 1μ*m*. The values used for the constants are: *D* = 30μ*m*^2^ ⋅ *min*^−1^, *C*_0_ = 1%, *C*_0_′ = 0.1%, *D*_*oxy*_ = 1300μ*m*^2^ ⋅ *s*^−1^, *r*_0_ = *ln* 2/480 *min*^−1^ and *b* = 0.1*O*_2_%*min*^−1^*cell*^−1^.

## Acknowledgments

We thank R. Fulcrand for his expert help in microfabrication; G. Torch, C. Zoude, G. Simon and A. Piednoir for technical assistance. This study was supported by the CNRS - Mission pour les Initiatives Transverses et Interdisciplinaires – « Défi Modélisation du vivant - 2019», by the GDR ImaBio (AMI fellowship) and by the IFS LyC Collaborative Research Project 2019 (Tohoku University). S. Hirose was supported by the STARMAJ Program (France-Japan: Research Internships for Master’s Students, Université de Lyon) and K. Funamoto by the CNRS (invited researcher position). This project has received funding from the European Research Council (ERC) under the European Union’s Horizon 2020 research and innovation programme (grant agreement No 639638 to V.Calvez).

## Author contributions

KF, VC, PG, IM, CA, JPR and OCE designed the research. PG, IM, BA, CA and JPR adapted the spot assays The microfluidic chambers were developed by JPR, KF and SH. Oxygen sensors were developed and manufactured by JPR, OCE, IM, KF and SH. OCE, BA, SH, KF, CA and JPR performed experiments. MD, VC and OCE performed numerical simulations and calculations. OCE, JPR, CA, MD and VC wrote the paper and all authors edited the manuscript.

