## Supplementary Information for "Hypoxia triggers collective aerotactic migration in *Dictyostelium discoideum*"

1. Supplementary experimental methods
   1.1. Cell growth and development
   1.2. Observations and analysis of self-generated aerotaxis by cell confinement (spot assay)
   1.3. Cell tracking, diffusion coefficients and aerotactic biases
   1.4. Microfluidic based oxygen gradient generator: design, fabrication and cell injection
   1.5. Gas control and injection
   1.6. Oxygen sensing film preparation
   1.7. Fluorescence microscopy for oxygen measurements
   1.8. Oxygen sensing film calibration
2. Supplementary numerical simulations
   2.1. Numerical simulation of oxygen tension
   2.2. Potts model simulations
3. Supplementary information
   3.1. Mathematical analysis of the 'Go or Grow' model
   3.2. A complementary viewpoint of the contribution of cell growth to the collective movement
4. Supplementary Table and Figures
5. Captions for Supplementary Movies
6. References

#### 1. Supplementary experimental methods

##### 1.1. Cell growth and development

Two cell lines were used: AX2 and their SodA+ mutant counterpart. The [Act15]:SodA strain (Bloomfield & Pears, 2003) was obtained from Dicty Stock Center (strain DBS0237194). Both cell lines were cultured similarly as follows. Cells were grown in HL5 media (Formedium, Norfolk, UK) at 22ºC with shaking at 180 rpm for oxygenation (Sussman, 1987). When appropriate, 20 µg/ml of G-418 was added to maintain selective pressure on overexpressing mutants. Exponentially growing cells were harvested, counted and cell density was adjusted at a given value for each type of experiment as indicated below.

##### 1.2. Observations and analysis of self-generated aerotaxis by cell confinement (spot assay)

For the spot assay, 1 µL of cell suspension containing 1000-8000 cells (typically 2000) was carefully deposited on a dry surface (the nunclon treated surface of Nunc 6 wells polystyrene plates (ThermoFisher Scientific, Waltham, MA, USA) or polydimethylsiloxane (PDMS, Sylgard 184, Dow Corning, Midland, MI, USA) for oxygen sensing) using a 1 µL syringe (Hamilton, Reno, NV, USA). The drop was incubated for 5 to 7 minutes in humid atmosphere at 22°C before gently adding 2 ml of HL5 medium without detaching the spotted cells forming a micro-colony. A 14-mm or 18-mm diameter round glass coverslip cleaned in ethanol, thoroughly rinsed in HL5

was kept wet and deposited on top of it. In some experiments, fluorescein FITC at 16µM was added to the HL5 medium and confocal slices were taken, showing that confined *Dictyostelium* cells were not compressed by the coverglass but separated from it by a layer of medium of about 50 µm (Fig. S1).
The outward spreading of the *Dictyostelium* micro-colony was observed at 22°C in transmission with three types of microscope: (i) a TE2000-E inverted microscope (Nikon, Tokyo, Japan) equipped with motorized stage, a 4x Plan Fluor objective lens (Nikon) and a Zyla camera (Andor, Belfast, Northern Ireland) using brightfield for most of the experiments lasting 16 h (Fig. 1A), (ii) a binocular MZ16 (Leica, Wetzlar, Germany) equipped with a TL3000 Ergo transmitted light base (Leica) operated in the one-sided darkfield illumination mode and a LC/DMC camera (Leica) for experiments over days (Fig. 1B) and finally (iii) a confocal microscope (Leica SP5) with a 10x objective lens for a few larger magnification experiments (Fig. 1C).
For computing densities, cell positions were determined using the built-in Find Maxima plugin in ImageJ (National Institutes of Health, Bethesda, MD, USA) through a custom made routine. Data analysis and plotting was performed in Matlab (Mathworks, Natick, MA, USA). For density profiles (Fig. 1E) and kemographs (Fig. 1D), the center of the colony was defined as the center of mass of all cells detected at all times. Cell positions were then turned into radial coordinates and cells were counted within concentric crown regions. Densities were calculated by dividing this count by the area of each crown.
Density profiles such as the ones showed in Fig 1E were treated to automatically extract the position, width and density of a ring in various experiments and at various time points. Density profiles were first stripped of values lower than 500 cell/mm$^2$ in order to avoid asymmetric baselines behind and in front of the ring. Resulting profiles were then fitted in Matlab by a Gaussian function with a non-zero baseline. The non-zero baseline corresponds to density in the bulk, the maximum of the Gaussian gives ring position, its height added to the non-zero baseline gives the cell density in the ring and its width the width of the ring.

##### 1.3. Cell tracking, diffusion coefficients and aerotactic biases

After retrieving cells' positions with optimized ImageJ macros based on Find Maxima, the individual trajectories were reconstructed with a squared-displacement minimization algorithm (http://site.physics.georgetown.edu/matlab/). Data were analysed using in house Matlab programs. Timelaspse microscopy experiments devoted to cell tracking in the spot assay experiments was acquired at a high frame rate (1 frame per 15s) (Figs. S4-S5) due to the very high cell density in the ring region (up to 2000 cells/mm$^2$, Figs. 1E, 3C). For the microfluidic experiments, as cells were plated at a lower density (less than 200 cells/mm$^2$), 1-min time intervals was used to track cell trajectories (Fig. 2C). In order to highlight aerotactic biaises, cells displacements over various time lags *dt* (*dt* up to 60 min) were projected in the radial direction for spot assays and in the gradient direction X for microfluidic experiments and eventually divided by *dt* to obtain velocity biases. Individual biases were then averaged within bins of equal distance (Figs. 2D, S5, S13). Individual effective cell diffusion constants were measured as the square of their displacement over their entire trajectory divided by the trajectory time length and divided by 4. These measurements were then similarly averaged over bins (Fig. S4).

##### 1.4. Microfluidic based oxygen gradient generator: design, fabrication and cell injection

A schematic of the present double-layer microfluidic device is shown in Fig. 2A. It is made of several layers of PDMS mounted on a bottom glass coverslip. The overall diameter *D* of the microfluidic device is 27 mm and the overall thickness *H* is 4 mm. Three parallel media channels are positioned for cell culture, and two gas channels are positioned at a height $H_g$=0.5 mm above the media channels to allow gas exchange between the channels during cell culture. The horizontal distance between the two gas channels narrows step-by-step (2 mm, 1 mm and 0.5 mm gaps, respectively), thus yielding to generate different gradients of oxygen concentration along the media channels simultaneously. All channels are 125 µm high and 2 mm wide, and therefore, the media and gas channels are separated by a PDMS wall of 375 µm thickness. A polycarbonate (PC) film (26 mm in diameter and 0.5 mm thickness) is embedded inside the device at a height *H*f=1 mm from the bottom coverslip to prevent oxygen diffusion from the atmosphere. The cartesian coordinate origin was set at the center of the media channel (median axis), and the *x* and *y*-directions were defined as parallel to media and gas channels respectively (Fig. 2A). The *z*-direction was set to the vertical direction from the top of the bottom coverslip.

The manufacturing steps are as follow. The media channel and gas channels were drawn with AutoCAD (Autodesk, Mill Valley, CA, USA) and replicated in SU-8 photoresist using classical photolithography techniques. These SU8 molds were silanized to make it non-adherent and reusable. PDMS was mixed at a 10:1 ratio of base:curing agent, poured over each mold to a thickness of $H_g$, and cured in an oven at 60 °C for more than four hours. On top of the cured PDMS layer of the gas channels, the above-mentioned PC film with 3 mm port holes punched at the locations of the media and gas channel ports was positioned. Additional PDMS was then poured over the PC film until the total PDMS layer became 3.5 mm thick, then the PDMS layer was cured in an oven at 60 °C overnight. The PDMS layers of the media and gas channel patterns were peeled off the silicon wafers and cut into 27 mm diameter circles. The PDMS layer with the gas channel pattern was punched to form inlets and outlets 2 mm in diameter to allow the infusion of gas mixtures. The channel-patterned surface of the PDMS layer with the gas channels and the top surface of the other PDMS layer with the media channels were plasma treated (PDC-001-HP; Harrick Plasma Inc., Ithaca, NY, USA) to bond with each other. After incubating the bonded PDMS mold overnight in an oven at 60 °C, 2 mm diameter inlets were punched to allow access to the media channels, respectively. Finally, the channel-patterned side of the PDMS mold and a 35 mm-diameter glass bottom dish with or without covered by an oxygen sensing film were plasma treated and bonded each other.

*Dictyostelium* cells were seeded in the media channels at density of $2x10^6$ cells/ml, and the cell culture medium was filled in the glass bottom dish up to the height covering the PDMS mold. Cells were allowed to adhere to the bottom surface (bare glass or coverglass covered with a sensing film) for 15min.

#### 1.5. Gas control and injection

We used a controlled oxygen concentration for three types of experiments: (i) to calibrate oxygen sensing films (see below), (ii) to create the oxygen gradients within microfluidic devices (see below) or to insure a pure hypoxic condition (pure $N_2$) at the end of the spot assay experiment. The gas mixture (0% to 21% $O_2$ in $N_2$) was prepared in a gas mixer (Oko-lab 2GF-MIXER to mix compressed AIR with 100% N2 or HORIBA STEC MU-3405, Kyoto, Japan to mix pure O2 and N2) by mixing pure $O_2$ (or air) and pure $N_2$. Free sensing films for calibration (i) or for the spot assay (iii) were placed inside 6-wells plates and the multiwells were placed in an environmental chamber fitting our microscope stage (H301-K-frame, Okolab , Pozzuoli, Italia).Gas was injected at about 500 mL/min in this chamber. Eventually, multi-wells were drilled to a diameter of 25 mm and the sensing films were glued with a silicone adhesive on the plate bottom to reduce the background noise from fluorescence. For microfluidic experiments, the tubes from the mixer were connected to the gas channels and gas was injected at a controlled flowrate (between 60 and 180 mL/min) into the device.

#### 1.6. Oxygen sensing film preparation

Oxygen Sensing films were prepared by inserting the luminescent O2 sensitive dye 5,10,15,20-Tetrakis-(2,3,4,5,6-pentafluorophenyl)-porphyrin-Pt(II) (PtTFPP, Por-Lab, Porphyrin-Laboratories, Scharbeutz, Germany) in a 4:1 PDMS:curing agent thin layer spin-coated on 30-mm to 35-mm rounded coverglasses. Briefly, 17 mg of

PtTFPP was dissolved in 5 mL chloroform and thoroughly mixed with 2.8 mg of PDMS and 0.7 mg curing agent. The mixture was degassed in a vacuum chamber for 5 hours. About 0.5 mL to 1 mL of this solution was spread on the coverglass and spin-coated for 2 min at 500 rpm with a final speed of 2000 rpm during 10s to flatten the edge bead. Chloroform was allowed to evaporate overnight while the polymer cured at 60°C. The final PtTFPP sensor film had a dye concentration of 4 mmol/L and was 25 µm thick. This thickness was measured using a ContourGT-K 3D Optical microscope (Bruker, Billerica, MA, USA) after removing a piece of film with a surgical blade. Sensing films were stored in dark. They were used to measure the oxygen concentration in self-generated $O_2$ gradients (spot assay) and for microfluidic experiments with controlled $O_2$ gradients.

#### 1.7. Fluorescence microscopy for oxygen measurements

Fluorescence images of $O_2$ sensing films (either for film calibration or for in situ oxygen measurements in the spot assay or in microfluidic devices) were taken with two inverted epifluorescence microscopes: (i) a TE2000-E inverted microscope (Nikon) equipped with motorized stage, a 4x Plan Fluor objective lens (Nikon), a X-Cite Series 120PC illumination lamp, a TRITC bandpass filter cube and a Zyla camera (Andor) (Fig. 3Aii, S8), (ii) a IX83 inverted microscope (Olympus, Tokyo, Japan) equipped with a motorized stage, a UPlanSApo 4x objective lens (Olympus), a U-HGLGPS lamp (Olympus), a RFP bandpass filter and a Zyla camera (Andor). This second microscope was used for mosaic imaging, in order to scan the whole dimension of the three media channels (about 1 cm in length) thanks to the dedicated imaging software cellSens (Olympus) (Fig. S6).

#### 1.8. Oxygen sensing film calibration

Calibration was carried out with the sensing films in air, in water or in HL5 culture medium. We applied gas concentration ramps with steps of 5 min for calibration in air (time to exchange fully the gas composition of the chamber and tubes, as $O_2$ almost instantaneously diffuses within the 25µm thick sensing film) and with much longer steps (*i.e.*, 2-4 h) for calibrations in liquid. There is indeed an additional diffusion time in the PDMS intermediate layer of our microfluidic devices or in the medium height of a Petri dish: typically a few minutes for a 0.5-mm thick PDMS layer and 1h30 for a 2.7-mm thick liquid layer in a dish.
Timelapse fluorescence images we recorded and signal intensity *I* was measured in ROIs of typically 64x64 pixels in various positions of the image, especially along a line scanning the middle of the image (Fig. S6B, S8A). The response of the sensing film in the presence of oxygen can be modeled by a linear Stern-Volmer relationship:

$$\frac{I_0 - B_g}{I(C) - B_g} = 1 + K\ C$$

where C is the oxygen concentration expressed as a percentage of oxygen in the injected gas phase (nearly 21% for atmospheric conditions), $I_0$ is the reference intensity in the absence of oxygen, $B_g$ is the background intensity independent from the oxygen sensitive signal of the PtTFPP molecules and *K* is the Stern-Volmer constant used as an indicator of the sensing film sensitivity.

Notice that the background is usually not included in the Stern-Volmer relationship but a representative background image ($O_2$ independent) is subtracted prior to intensity measurements (Nock, Blaikie, & David, 2008; Thomas et al., 2009). This $O_2$ independent background value can come the fluorescence of a plain PDMS film prepared in the same conditions than the sensing film but devoid of PtTFPP molecules (*i.e.*, a standard) (Thomas et al., 2009). We tested that procedure that is basically working but we choose to include the background as a fitting parameter because illumination conditions may change between the sample and the standard (especially the focus plane that affects the focused height of autofluorescent medium above the surface). The slightly varying thickness and PtTFPP composition of sensing films at the large spatial scales we are interested here (3 to 6 mm wide images, Figs. S6A-D, S8A-C) are another sources of heterogeneity especially for the *K* value. For those reasons, we apply the Stern-Volmer relation with *K* and $B_g$ as a fitting parameters in many different small regions of interest (ROI) of the surface. For each ROI, we found that the measured intensities follow perfectly the Stern–Volmer relation (*i.e.*, $C$ linearly increases with the Stern-Volmer parameter $(I_0\text{-}B_g)/(\ I(C)\text{-}B_g)\text{-}1$, Fig. S7B) and that $K$ and $B_g$ are clearly uncorrelated, in particular $B_g$ depends on the illumination pattern but not $K$.

The illumination pattern is clearly visible on fluorescence images at 21% $O_2$. For instance, the large field of view of Fig. S6B reconstituted by the multi-area module of the microscope displays an up and down landscape in the 21% intensity (Fig. S7C) due illumination changes in the periphery of each overlapped area but also due to the slightly different fluorescence in the gap region of the microfluidic device and especially at its interfaces. The single large field of view of Fig. S8A taken with our second microscope setup displays a dome shaped pattern with 20% intensity difference between the center and borders (Fig. S8D). The background value $B_g$ is very correlated with the 21% signal variations in both imaging configurations (Figs. S7C, S8D) and we can for each experiment calibrate a linear relationship between between $B_g$ and $I(21\%)$ (Figs. S7D,S8F).

The background is due to the autofluorescence of the glass supporting the sensing film, of the PDMS chains constituting the matrix of the 25-µm thick sensing film and of the fluid surrounding it. For Fig. S6, the microfluidic device was filled with pure water and for Fig. S8, the calibration was performed in the autofluorescent HL5 medium before spotting the cells. Sometimes the sensing film was placed in a non drilled well and in that case the strong autofluorescence of the plastic bottom of the plate becomes a major source of background (not shown). Finally, $B_g$ also includes the read noise $RN$ of the camera which is a constant independent of the light output or exposure time. For Fig. S7, we measured $RN$=108 A.U. and hence a light background $B_g^*= B_g\text{-}RN$=15 A.U. which is half the "true oxygen dependent signal" at 21% $O_2$, $I(21\%)\text{-}RN$=30 A.U. The maximum deviation of $B_g$ from the linear fit in Fig. 10D is about 1.5 A.U. Hence a relative error 1.5/15=10% for $B_g^*$ will be taken in the following. For Fig. S8, we measured $RN$=100 A.U. and at the top of the bell curve, $B_g^*$=1400-100=1300 A.U. while $I(21\%)^*$=1600-100=1500 A.U. (Fig. S8D). Hence, the background is nearly 87% of the signal due to the HL5 autofluorescence. Nevertheless, the maximum relative deviation from the linear fit is smaller at about 25/1300≈2 % of $B_g^*$. All these values will be used for the error analysis of the oxygen profiles below.

For uncovered (free) sensing films he sensibility $K$ ranges between 3 to 5 $\%^{-1}$ and is very constant, weakly dependent on the illumination pattern (Fig. S7D and blue points in Fig. S8E). When films are covered with a coverglass, the fluorescence under hypoxic conditions (0%) increases significantly on the covered region (Fig. S8C) but not at 21% (Fig. S8B). As a results, K, which is proportional to this ratio, increases significantly (Fig. S8E) but not $B_g$ and $I$(21%) (Fig. S8D) confirming that $K$ and $B_g$ are independent. This increase in $K$ is probably due to a local temperature increase: the coverglass adsorbs more heat from the light and this heat is difficult to evacuate due to the confinement. In principle, for the spot assay experiment, it would be necessary to perform an independent calibration with the Stern-Volmer relation in the covered situation. However, this is difficult due to the very long time required to equilibrate the oxygen level under the confinement far from the coverglass boundary (this is why we started the Stern-Volmer fit at $ROI_7$ in Fig. S8B,D,E). A too long procedure causes other problems such as medium evaporation, stage or focus drift… To avoid that, we decided to apply the protocol described in the image analysis pipeline of Fig. S9. First, we perform a gas calibration ramp and do a Stern-Volmer analysis in various points of an uncovered sensing film (the subscript $U$ is for uncovered) where $I_{0U}$ is reliable in order to get the linear background relation $B_{gU}=\alpha\, I(21\%)_U+\beta$ and to measure the measure the ratio $R=I_{0U}/\mathrm{I}(21\%)_U$. Reliable means here any point if gas mixture is applied uniformly when calibrating in a dish or just underneath the gas channel in microfluidic devices. Second, we choose the reference fluorescence image $I(21\%)$ immediately before starting any experiment (*i.e.*, just after covering the spot, or just before applying the gradient in the gas channels). From that image, we build a $B_g$ image as $\alpha\, I(21\%)+\beta$ (as $B_g$ is the same for uncovered and covered case) and eventually we build a reconstituted $I_0$ image as $R\, I(21\%)$. Finally, we subtract and divide images with the Image calculator of ImageJ (*i.e.*, pixel by pixel) following the Stern-Volmer model and hence get a $K$-value image map and subsequently an oxygen map (Fig. S9C-D). This enables to correct non-homogeneous illumination conditions or non-homogeneous sensing film properties as well to quickly estimate error bars on the oxygen map from the estimated errors on $B_g$ , *I(21%)* and $I_0$ detailed below.

We already discussed the error on the background. In principle, *I(21%)* is a reference image (hence error free), however as an experiment (especially the spot assay) may run overnight we need somewhere to evaluate drift in the absolute intensity for instance by running an overnight timelapse experiment with a sensing film under ambient gas conditions. This error is added on *I(21%)* and was estimated as 2% of the true fluorescence signal corrected from read noise *I(21%)-RN*. Error on $I_0$ could be much larger. As the intensity *I(C)* is strongly nonlinearly increasing with the oxygen concentration *C,* if we measure an intensity $I_0*$ corresponding to a residual small oxygen level $C_0*$, we need to correct the true $I_0$ value using the relation $I_0 - B_g \approx I_0 \approx I_0^{*}(1 + K\,C_0^{*})$. Due to oxygen leakage along tubes and within our environmental chamber, when applying 100% $N_2$, we measured a residual $C_0^{*} \approx 0.15\%$ $O_2$ in a culture medium dish using a bare fiber oxygen sensor coupled to its commercial oxymeter (Firesting, Pyroscience, Aachen, Germany). Hence with a typical *K*=5 value, we obtain a very large discrepancy between the measured fluorescence $I_0*$ and the ideal one: $I_0 \approx 1.75\, I_0^{*}$, but finally this discrepancy is not really dramatic on the measured error again due to the non-linearity.

The effect of these different error sources on the measured oxygen map is presented in Figs. S6E and S8H for a typical microfluidic and spot experiments. Even if we make a 1.75 error on $I_0$, this has little effect on the profiles except in the very hypoxic region when C<0.25% where the error exceed 50%. But even in the region around C=1%, the error is less than 10%. The error on *I(21%)* on the other hand has a significant effect on the high oxygen regions but less on the hypoxic regions. Finally, the background error is relatively visible in the intermediate oxygen concentration region (very visible on the side of the spot in Fig. S8H, but also to some extend around the median axis at C~10% in the microfluidic experiment, Fig. S6E). Overall, we defined error bars with min-max values of this bootstrap error analysis. This error in Fig. 2B of the article. In the 0.5-1.5% region were we observe most of the interesting aerotactic behaviors with *Dictyostelium* cells, the precision on the oxygen concentration is better than 30%. But this absolute error is probably the same for a same device whatever the location. The relative error between the three gaps or between different locations along the gradient axis are likely to be much smaller. For the purpose of this paper, we can conclude that aerotaxis and aerokinesis occurs undoubtedly at oxygen concentrations between 0% and 2% (Fig. 2).

#### 2. Supplementary numerical simulations

##### 2.1. Numerical simulation of oxygen tension

Oxygen tension inside the device was computed using commercial finite element software (COMSOL Multiphysics 5.5; COMSOL, Inc., Burlington, MA, USA). The gas flow in the individual channels were simulated by solving the Navier-Stokes equations coupled with mass continuity for an incompressible fluid:

$$\rho_G(\mathbf{u}\cdot\nabla)\mathbf{u} = \mu_G\Delta\mathbf{u} - \nabla p,$$
$$\rho_G\nabla\cdot\mathbf{u} = 0,$$

where **u** is the velocity vector, *p* is the pressure, and $\boldsymbol{\rho_G}$ and $\boldsymbol{\mu_G}$ are the gas density and viscosity (taken as 1 kg/m$^3$ and 10$^{-5}$ Pa.s respectively). The spatial and temporal distribution of oxygen inside the device was then calculated by solving the convection-diffusion equation:

$$\frac{\partial c}{\partial t} = D\Delta c - \mathbf{u}\cdot\nabla c,$$

where *c* is the oxygen concentration, *D* is the diffusion coefficient of oxygen, and *t* is the time.

The device was assumed to be in an atmosphere containing 21% $O_2$. Medium at 21% $O_2$ concentration was supplied to media channels. Gases containing 0% and 21% $O_2$ were respectively supplied to the left-hand and right-hand side gas channels at 30 ml/min to generate an oxygen gradient.

Zero pressure and convection flux conditions were set at the outlets of the gas channels, and a no-slip condition was applied on the channel walls for fluid flow analysis. Boundary conditions for oxygen concentration were set according to Henry's law. Oxygen concentration at the interfaces between the PDMS and gas phase (atmosphere and gas mixture in the gas channel) was set correspondingly to the product of the solubility coefficient of oxygen in PDMS and the partial pressure of oxygen. At the interfaces between PDMS and media or gel, a partition condition was applied, which balanced the mass flux of oxygen to satisfy continuity of partial pressure of oxygen:

$$\frac{c_{\mathrm{PDMS}}}{S_{\mathrm{PDMS}}} = \frac{c_{\mathrm{channel}}}{S_{\mathrm{channel}}},$$

where $c_{\mathrm{PDMS}}$ and $S_{\mathrm{PDMS}}$ are the oxygen concentration and the solubility of oxygen in the PDMS, respectively, and $c_{\mathrm{channel}}$ and $S_{\mathrm{channel}}$ are those in the media and gel channels. Moreover, oxygen consumption by cells was considered by setting an outward flux of oxygen of $6\times10^{-8}$ [mole/($m^2$.s)] on the bottom of the media channels (calculated as $b\,\rho^*$ where $b=1.2.10^{-16}$ mole/(cell.s) is the oxygen molar consumption per *Dd* cell per unit of time (Torija et al., 2006) and $\rho^*=500$ cell/$mm^2$ is the highest density used in the device). The initial condition of oxygen concentration in each material was set to 21% $O_2$. The physical properties of materials such as density, viscosity, oxygen diffusivity, and oxygen solubility for the medium, gas, PDMS, and PC film are summarized in Table ST1. The solubilities of oxygen in PDMS and the PC film were assumed to be the same since they are reportedly within the same range (Merkel, Bondar, Nagai, Freeman, & Pinnau, 2000; Moon, Monson, & Extrand, 2009). The computational models consisted of approximately 1,135,000 computational elements.

**2.2. Potts model simulations**

In this section, we will discuss the details of the Potts model. As presented in the main text, some parameters were directly taken from experimental observations and some were fitted by trial and error to faithfully reproduce the microfluidics experiments.

In all simulations, we used Compucell's Volume module which applies to all cells a Hamiltonian of the form:

$$H_{volume} = \lambda_v (V - V_{cell})$$

Where $V$ is the volume of a cell and $V_{cell}$ a target volume set to 2 pixels. This already set the length scale of our simulations to 1pixel = 10µm. $\lambda_v$ was set to 800. These values were adapted to reproduce the cell speeds observed in the microfluidic experiments. To achieve this relationship, we also decided to fix that one step of the simulation (Monte Carlo Step) was meant to represent 0.1s.

Aerotaxis was modeled using CompuCell's built-in chemotaxis plugin. This leads to a new term in the Hamiltonian of the form

$$H_{chemotaxis} = \lambda_{aero} \Delta C$$

Where $\Delta C$ is the difference in oxygen concentration $C$ between the source and target pixels of a flip and $\lambda_{aero}$ is the aerotactic strength. Key to our model is thus the fact that we made $\lambda_{aero}$ different for each cell and dependent on the local oxygen concentration. This modulation was fitted to the microfluidic experiments and set, in the general model as:

$$\lambda_{aero}(C) = \frac{800}{1 + e^{C-0.7/_{0.2}}}$$

Where C is the oxygen concertation at the center of mass of a cell. Fig 5 shows variations on that relationship which are:

$$\lambda_{aero}(C) = \frac{1225}{1+e^{C-0.7/_{1.5}}};\ \lambda_{aero}(C) = \frac{1375}{1+e^{C-0.7/_{3}}} \text{ and } \lambda_{aero}(C) = 700$$

Based on experimental observations, we also made the effective temperature of the model different for each cell and dependent on local oxygen concentrations. This allowed us to reproduce the aerokinetic effect and the modulation of the temperature was fitted to reproduce the cell diffusion constants measured in the microfluidics experiment. The main model thus uses the following relationship for temperature T

$$T(C) = 85 + \frac{105}{1 + e^{C-0.7/_{1}}}$$

Fig 5 and Fig S16 show variations on this relationship which is simply replaced by a constant value of T (115, 135 or 155).

Another key aspect of the models is the oxygen field which is implemented using CompuCell's DiffusionSolverFE module. In the case of the microfluidics experiments, the oxygen field was made to be constant (no diffusion, no consumption by cells) and fitted on experimental measurements of this gradient shown in Fig 2B. The oxygen concentrations were expressed in % giving 0 and 21 as natural boundaries. The actual oxygen profile varied in the x direction only as:

$$C(x) = \frac{21.38}{1 + e^{0.031*(x-200)}}$$

Where x is the position in pixels in the simulation which was run on a 400 by 800 grid (x and y respectively).

For spot assays, oxygen was allowed to diffuse freely. The diffusion constant of oxygen in liquids is on the order of $2.10^{3}$ $\mu m^{2}.s^{-1}$ (see Table ST1). In our time and length units, this yields a diffusion constant of 2 $pixel^{2}$ $step^{-1}$.

In terms of consumption, we took the oxygen consumption by Dicty cells to be $1.2.10^{-16}$ mole/(cell.s) (Torija et al., 2006). Given a solubility of oxygen in the medium of $2.5.10^{-4}$ mol $L^{-1}$ (measured with a bare fiber sensor plugged to a a Firesting oximeter, Pyroscience) and a vertical confinement of 50µm, the amount of oxygen available, at maximum, above a single pixel of the simulation is $1.25.10^{-15}$ mol which we define, in arbitrary units, to be 21. We can then turn the consumption of a single cell into a consumption per pixel given that the typical size of a cell is 2 pixels and per time step, each representing 0.1s. We end up with a consumption, in our arbitrary units of 0.1 $pixel^{-1}$ $step^{-1}$ which is only applied to pixels occupied by a cell. Of note, in case an occupied pixel had a remaining oxygen level of less than this values, then consumption was set at this oxygen level so that all oxygen was consumed. The last ingredient in oxygen dynamics is the leak of oxygen coming from the bottom of the multiwall plate. Assuming complete hypoxia on the cells' side, this would lead to a net flux of oxygen of *DC/e* where D is the oxygen diffusion constant in polystyrene, C is the oxygen concentration on the outside and e is the thickness of the polystyrene bottom. This leads to a flux by unit surface, in our Potts units of 0.001 $pixel^{-1}.step^{-1}$. We therefore implement a source of oxygen for all pixels in the simulation, whether they are occupied by a cell or not, of the form:

$$secretion\ (C) = 0.001\ \frac{21 - C}{C}$$

Where C is the local oxygen concentration at the considered pixel.

This was sufficient to faithfully reproduce the formation time of the rings. Finally, the spot simulations were run on a 500 by 500 pixels grid and we imposed boundary conditions to the oxygen field as a constant concentration of 21, the borders acting as a source of oxygen just like the edges of the coverslip in the experiments.

Cell division was also set to experimental observations. Given a doubling time of 8h, we implemented random divisions at each time point, each cell having a 1/(8h * 3600 s/h * 10step/s)= $3.10^{-6}$ chance of dividing. In Fig5, a simulation is shown were this probability was set to 0.

In terms of initial conditions, the microfluidic simulations were started from a homogenous cell density, each cell being initialized on a grid: 2 pixels per cells and a 6 pixel gap to the next neighbor in all directions. For the spot simulations, cells were seeded in three circular, concentric regions of decreasing density. The first region was set to be 30 pixels (300 µm) in radius with a gap of 1 pixel between each cell, the second one spanned the radii between 30 and 60 pixels with a gap of two pixels between each cell and the last one spanned between

60 and 90 pixels with a gap of 3 pixels. This lead to an initial colony with a radius of 900μm and between 1900 and 2000 cells, both very similar to experiments.

### 3. Supplementary information

##### 3.1. Mathematical analysis of the 'Go or Grow' model

We present below a preliminary analysis of the 'Go or Grow' model. A more detailed mathematical investigation of this model will be carried out in a separate article.

1. The 'Go or Grow' model admits explicit traveling wave solutions.
We recall that $z = x - \sigma t$is the spatial variable in the moving frame at (unknown) speed $\sigma > 0$. We seek a pair of stationary profiles, resp. the density $\rho(z)$and the oxygen concentration $C(z)$. We assume that $C(z)$is an increasing function. By translation invariance, we set without loss of generality that $C(0) = C_0$, so that EQ3 becomes:
[EQSI1]

$$\{ \begin{matrix} -\sigma \frac{\partial \rho}{\partial z} = D \frac{\partial^2 \rho}{\partial z^2} - a_0 \frac{\partial \rho}{\partial z} if z < 0 \\ -\sigma \frac{\partial \rho}{\partial z} = D \frac{\partial^2 \rho}{\partial z^2} + r_0 \rho if z > 0 \end{matrix} \}$$

The function $\rho(z)$must furthermore satisfy at $z = 0$ the following jump relation (ie. the continuity of the flux).

[EQSI2] $$\frac{\partial \rho}{\partial z}(0^+) - \frac{\partial \rho}{\partial z}(0^-) = \frac{-a_0}{D}\rho(0).$$

Thus the equation becomes a second order differential equation with piecewise constant coefficients on each half-line, that can be solved explicitly.
For $z < 0$ , the solution is of the form $A + Be^{\frac{a_0-\sigma}{D}z}$. From EQ3bis we observe that $\sigma \geq a_0$ equivalently $\frac{a_0-\sigma}{D} \leq 0$, which implies that $B = 0$.
For $z > 0$, we look at the roots of the characteristic polynomial $P(\mu) = D\mu^2 + \sigma\mu + r$. We note that to yield a nonnegative solution, we need $\sigma^2 \geq 4Dr_0$.
If $\sigma = 2\sqrt{Dr_0}$, then the solution is of the form $(Cz + D)e^{-\sqrt{\frac{r_0}{D}}z}$ and with the jump relation EQSI2, we obtain $\rho(z) = A(\frac{\sqrt{Dr_0}-a_0}{D}z + 1)e^{-\sqrt{\frac{r_0}{D}}z}$ and we observe that in this case, we necessarily have $a_0 \leq \sqrt{Dr_0}$.
If $\sigma > 2\sqrt{Dr_0}$, the solution is of the form $A'e^{\mu_- z} + B'e^{\mu_+ z}$, with $\mu_\pm = \frac{-\sigma \pm \sqrt{\sigma^2-4Dr_0}}{2D}$. By arguments exposed in (Van Saarloos, 2003), solutions with initial datum localized cannot decrease exponentially at a rate $\mu > -\sqrt{\frac{r_0}{D}}$, which leads to $B' = 0$, as $\mu_+ > -\sqrt{\frac{r_0}{D}}$. Then $A = A'$, but in order to satisfy the $C^1$-discontinuity jump relation [EQSI1], it must be that $\mu_- = \frac{-a_0}{D}$ [EQSI3].
[EQSI3] can algebraically be solved for $\sigma$ which yields $\sigma = a_0 + \frac{Dr_0}{a_0}$. Furthermore we can rewrite [EQSI3] as follows $(2a_0 - \sigma) = \sqrt{\sigma^2 - 4Dr_0}$, multiplying by $2a_0 + \sigma$, we find that $(4a_0^2 - \sigma^2) = (2a_0 + \sigma)\sqrt{\sigma^2 - 4Dr_0} > 0$, which leads to $a_0^2 > \frac{\sigma^2}{4} > Dr_0$.

Thus we have disclosed all possible profiles. In the case $a_0 \leq \sqrt{Dr_0}$ the only possible profile travels at speed $\sigma = 2\sqrt{Dr_0}$, whilst for $a_0 > \sqrt{Dr_0}$ the only possible profile travels at speed $\sigma = a_0 + \frac{Dr_0}{a_0}$.
One needs to verify that each of these profiles admits an associated oxygen profile that satisfies the condition $C(0) = C_0$, but the preceding profiles were defined up to the multiplicative constant $A$, by linearity of EQ3. The differential equation on $C$ becomes with $\tilde{\rho}$ the solution given above for $A = 1$:
[EQSI4]

$$-\sigma \frac{\partial C}{\partial z} = D_{oxy} \frac{\partial^2}{\partial z^2} - b(A\tilde{\rho})C$$

One concludes by checking that there exists a unique constant $A$ such that the solution to the differential equation satisfies $C(0) = C_0$.

2. The wave is pushed in the case $a_0 > \sqrt{Dr_0}$.
A neutral fraction $v^k$is defined as satisfying the following linear equation in the moving frame $z = x - \sigma t$:
[EQSI5]

$$\frac{\partial v^k}{\partial t} + Lv^k := \frac{\partial v^k}{\partial t} - \sigma \frac{\partial v^k}{\partial z} - D\frac{\partial^2 v^k}{\partial z^2} + \frac{\partial}{\partial z}(a(z)v^k) - r(z)v^k = 0, \text{ with } v^k(0,z) = v_0^k(z).$$

where we identify $a(z) = a(C(z))$and $r(z) = r(C(z))$ for the sake of clarity. This corresponds biologically to staining the cells given by the initial distribution $v_0^k$at time $t = 0$ with a neutral label (Roques, Garnier, Hamel, & Klein, 2012).
Defining $U(z) := \frac{(\sigma - a(z))}{D} z$, then we note that $Lf = -D\frac{\partial}{\partial z}(e^{-U}\frac{\partial}{\partial z}(e^U f)) - r(z)f$. This leads to setting $w := e^{\frac{U}{2}}v^k$that satisfies the parabolic equation $\frac{\partial w}{\partial t} + \tilde{L}w = 0$, with $\tilde{L}g := -De^{\frac{U}{2}}\frac{\partial}{\partial z}(e^{-U}\frac{\partial}{\partial z}(e^{\frac{U}{2}}g)) - r(z)g = -D\frac{\partial^2 g}{\partial z^2} + (\frac{U'^2}{4} - r(z) - \frac{a_0}{2}\delta)g$. The operator $\tilde{L}$ is self-adjoint in $L^2(\mathbb{R}, dz)$on the appropriate domain. Then by setting $\gamma := inf\{\frac{U'^2}{4} - r(z)\} > 0$, one can first show that every element of the spectrum of $\lambda \in \sigma(\tilde{L})$ such that $\lambda < \gamma$ is an eigenvalue of $\tilde{L}$. Second, one shows that the only eigenvalue $\lambda$ of $\tilde{L}$ such that $\lambda < \gamma$ is $\lambda = 0$. Finally by standard theory of self-adjoint operators and semi-group theory, one obtains that $w(t) = Pw_0 + e^{-t\tilde{L}}(I - P)w_0$, where $\|e^{-t\tilde{L}}(I-P)w_0\|_{L^2(\mathbb{R},dz)} \leqslant e^{-\gamma t}\|w_0\|_{L^2(\mathbb{R},dz)}$. Translating these properties onto the neutral fraction $v^k$, we have that $v^k(t) \to \frac{\langle v_0^k|\rho\rangle_{L^2(\mathbb{R},e^U dz)}}{\langle \rho|\rho\rangle_{L^2(\mathbb{R},e^U dz)}}\rho$ at an exponential rate, where $\rho$ is the traveling wave profile calculated in the previous section. Therefore each fraction converges to a fixed proportion of the whole population. We conclude that after some time the wave becomes a perfect mix of each neutral fraction. This corresponds to the definition of a pushed wave according to (Roques et al., 2012).

3. The wave is pulled in the case $a_0 \leq \sqrt{Dr_0}$.
A rigorous proof of this claim is still under investigation. The preceding reasoning does not apply to this case, as there seems to be no appropriate $L^2$-space where the methodology would apply. However, the intuition is clear, as the wave speed coincides with Fisher's $\sigma = 2\sqrt{Dr_0}$. This is the signature of a pulled reaction- diffusion front that does not involve the advection contribution at speed $a_0$. This intuition is confirmed numerically. We observed that the neutral fraction left to the peak vanishes in the moving frame. Additionally the rate of decay is not exponential, but rather a rate of order $\frac{1}{\sqrt{t}}$ (Fig S19). This excludes the possibility of being in the same framework as in the previous section 2. Furthermore, the results seem to be in good agreement with the analogous property for the Fisher-KPP case (Roques et al., 2012). It is shown that for the Fisher-KPP model any neutral fraction $z = x - \sigma t$ with initial datum $\int_{\mathbb{R}} e^{2\sqrt{\frac{r_0}{D}}x}(v_0^k(x))^2 dx < +\infty$, meaning that initially the

presence of the neutral fraction $z = x - \sigma t$ in the front is negligible, vanishes on sets of the form $[A, +\infty)$ with a rate of order $\frac{1}{\sqrt{t}}$. The different dynamics for pushed and pulled waves in the case of the simple 'Go or Grow' model are summarized in Fig S20.

**3.2 A complementary viewpoint of the contribution of cell growth to the collective movement.**

For the 'Go or grow' model or one of its variations, there exists an alternative viewpoint that combines the shape of the density profile and the wave speed. By integrating [EQ3] over the line, we obtain

[EQ3bis] $$\left(\sigma - a\big(C(-\infty)\big)\right)\rho(-\infty) = \int r\big(C(z)\big)\rho(z)dz$$

This equation balances the net flux of cells to the far left-hand with the amount of mass created by heterogeneous (signal-dependent) cell division.

Formula [EQ3bis] remains true under variations of the simple 'Go or Grow' model, as it merely results from mass balance. We illustrate this relationship with the experimental data from Fig. 1E. We approximate the area under the curve by a rectangle method: $\int r\big(C(z)\big)\rho(z)dz = r_0\hat{\rho}L$ where $L$ is the length spanned by the ring and $\hat{\rho}$ is the average cell density in the ring. As there is supposedly no advection $a(C) = 0$ at $z = -\infty$ this yields the approximation $\sigma \approx \frac{r_0 L\hat{\rho}}{\rho(-\infty)}$. Quantitatively, we assume L to be on the order of 300µm (Fig. 1E) and $\frac{\hat{\rho}}{\rho(-\infty)}$, the ratio between cell densities in the ring and in the bulk of cells, to be on the order of 4 (Fig. 1E). This yields an estimate of the wave speed of $\sigma \approx 1.7 \mu m.min^{-1}$ relatively similar to the estimate given by [EQ4].

### 4. Supplementary tables and Figures

| | Culture media | PDMS | PC (polycarbonate) |
|---|---|---|---|
| Diffusion constant D [$m^2/s$] | $2\ 10^{-9}$ | $4.1\ 10^{-9}$ | $2\ 10^{-12}$ |
| Solubility S at 1 atm. (µM) | 219 | 1666 | 1666 |

**Table ST1.** Parameters used for the COMSOL simulations of oxygen equilibrium profile in microfluidic devices (oxygen diffusion constant and oxygen solubility in the different materials crossed). From (Hamon, Hanada, Fujii, & Sakaif, 2012; Merkel et al., 2000; Moon et al., 2009).

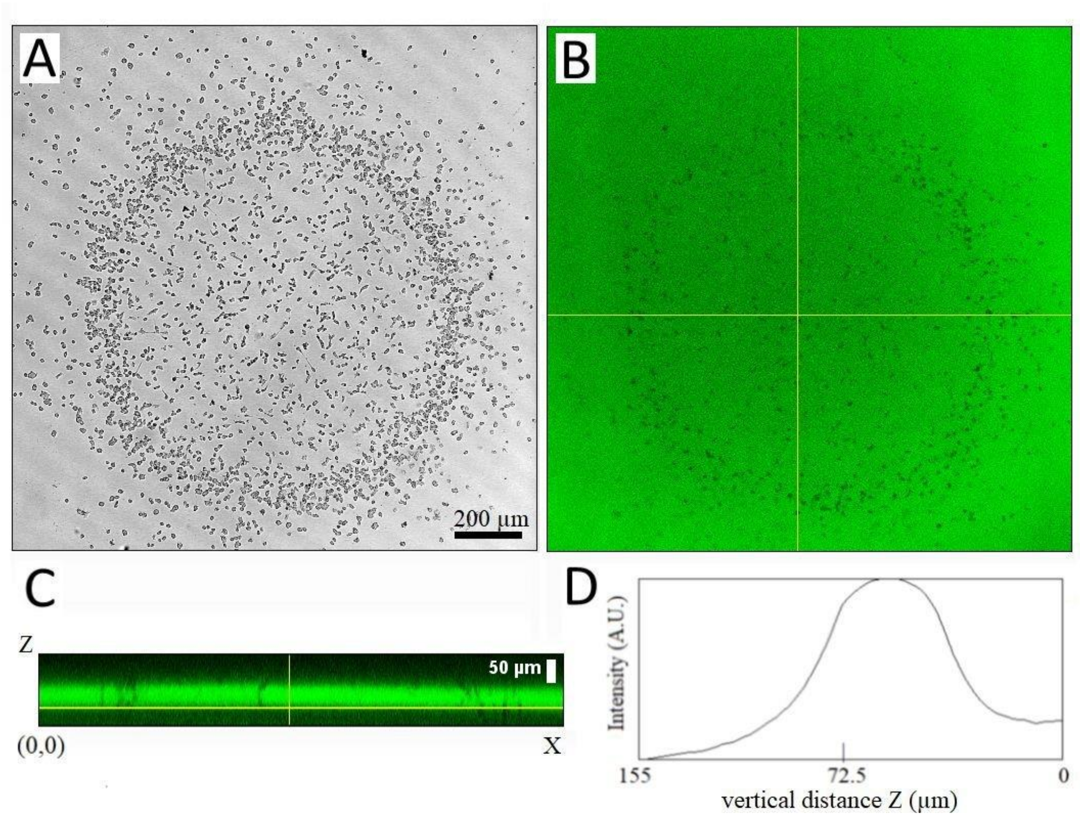


**Fig. S1.** Measurement of the confinement height 105 min after covering a cell colony with ~1000 cells plated on plastic with a coverglass using A-B) Confocal transmission and fluorescence XY images (slice 37) of a Z-stack from inside the plastic bottom of the well to the coverglass; C) Side view XZ along the horizontal line in B). D) Vertical intensity profile along Z (averaged over X). The Full width at half maximum (FWHM) is about 50µm. Z-stacks were taken with a confocal microscope at 2.57µm/slice using a 10x objective with a very small pinhole size (0.25 Airy), the fluorescence is due to fluorescein FITC added at 16µM in HL5 medium.

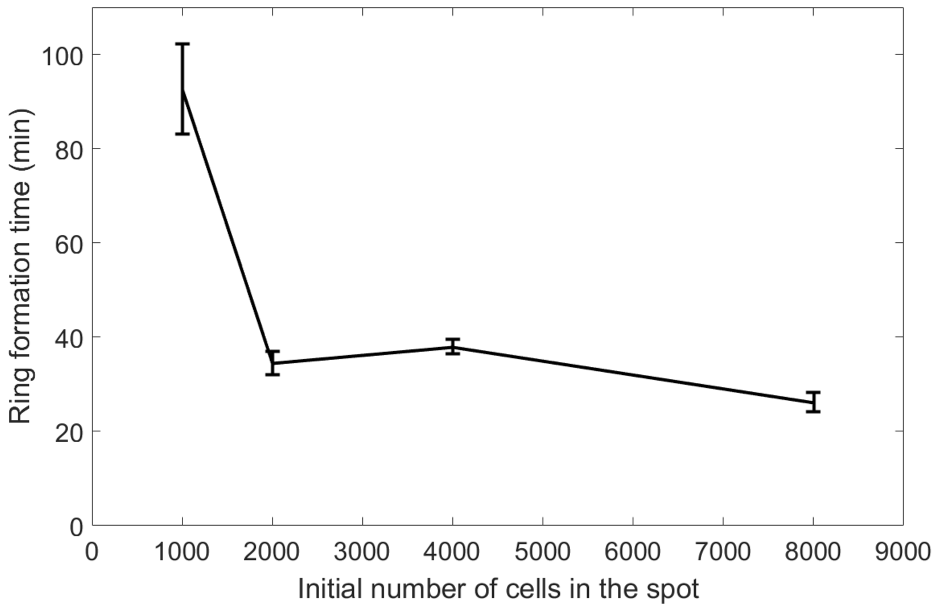


**Fig S2**. Ring formation time decreases with cell number. Error bars represent std of n>=4.

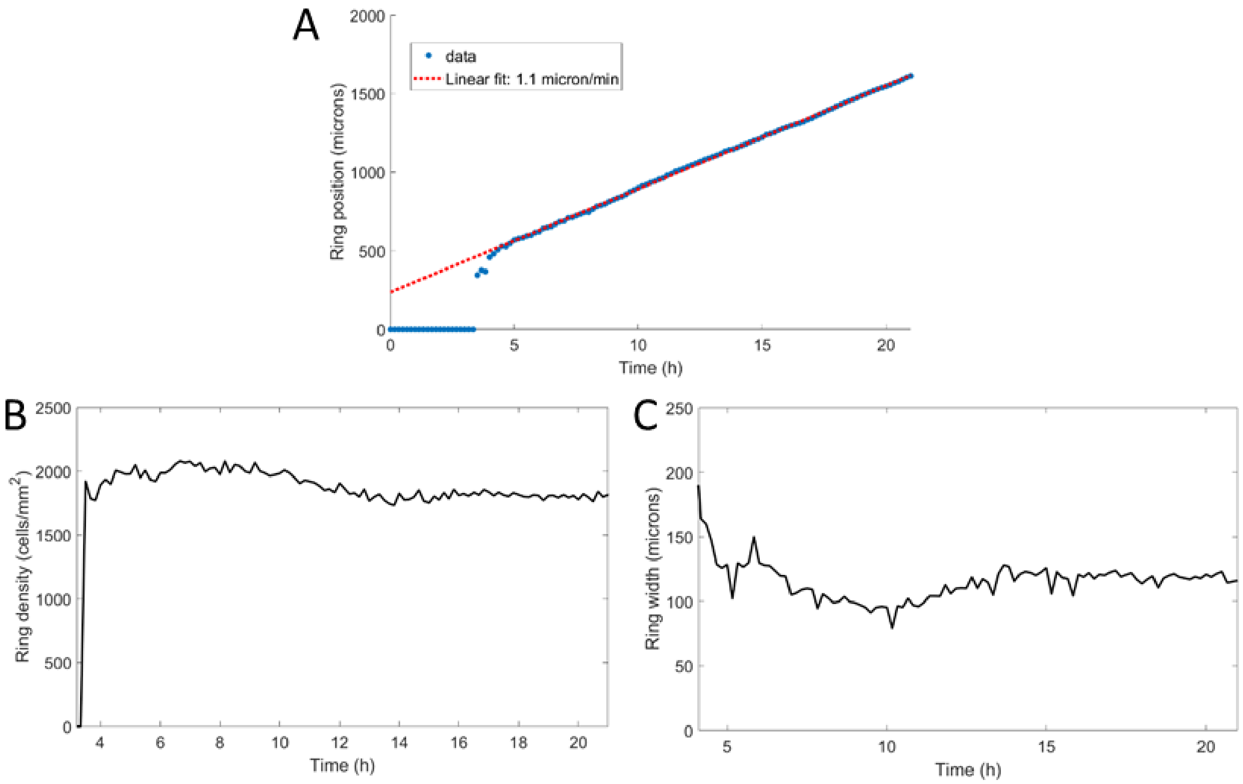


**Fig S3**. A*: Position of the ring as a function of time with a linear fit yielding a speed of 1.1μm/min. B: Cell density within the ring as a function of time, starting after ring formation. C: Ring width at half height as a function of time, after ring formation.*

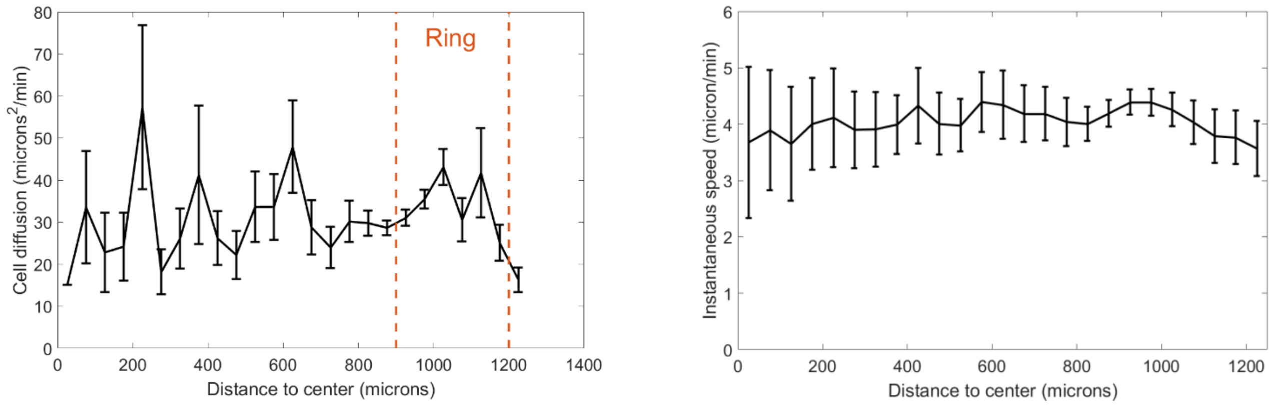


**Fig S4.** Effective cell diffusion constant and instantaneous speeds as a function of distance to the center. Cell diffusion is defined as the squared of the migrated distance during a trajectory divided by 4t with t the duration of that trajectory. Cells were tracked between t=9h and t=10h and orange dashed line represent the position of the ring at these times. Each point is the mean±std of cells within the given distance bin, n=2746 cells total.

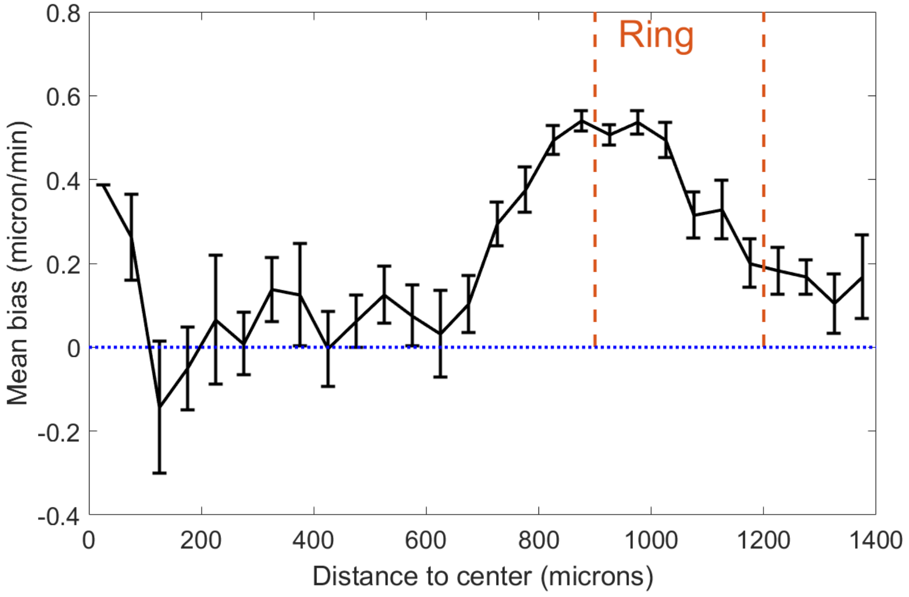


**Fig S5.** Cell velocity bias in the spot assay as a function of distance to the center. This bias is defined as the projected speed in the radial direction. Each point is the mean±std of cells within the given distance bin, n=2211 cells total. Cells were tracked between t=9h and t=11h and orange dashed lines represent the positions occupied by the ring at these times. Dotted blue line is a guide for the eye representing a 0 bias, *i.e.* non-oriented motion.

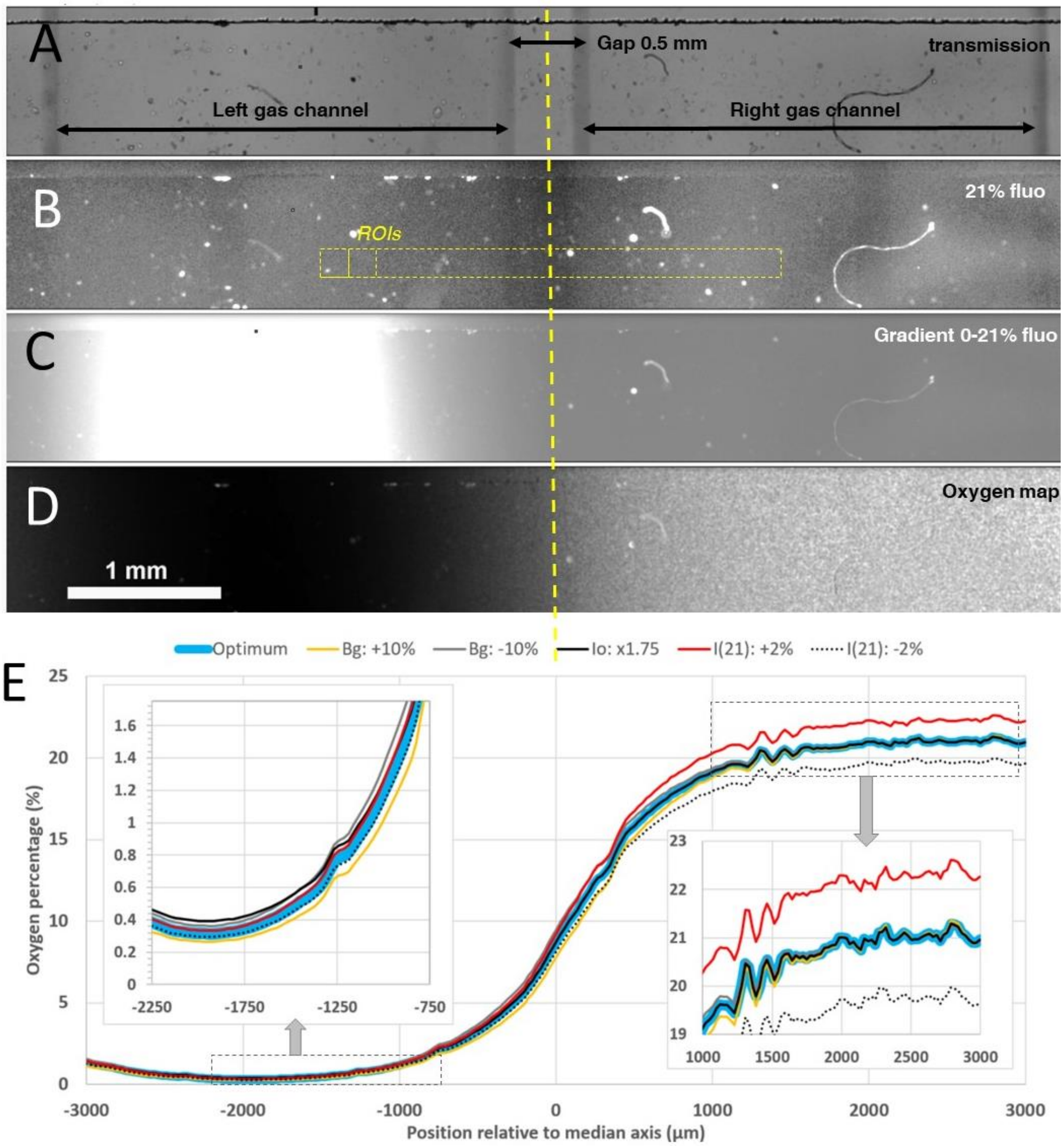


**Fig. S6.** Oxygen profile measurements inside the microfluidic gradient generator device with a sensing film mounted on the bottom of the media channel. A-B) Transmission and homogeneous 21% $O_2$ level fluorescence images of the media channel with gap 0.5 mm between the two gas channels. C) Raw fluorescence image in presence of 0-21% gradient established by injecting pure N2 in the left gas channel and air in the right one (image was taken 30 min after the beginning of the gas injection). D-E) Corresponding calculated $O_2$ map and $O_2$ profile. In D) colors correspond to slight changes within the experimental uncertainty of the parameters used for the calculation (see text), insets correspond to the hypoxic (~0%) and normoxic (~21%) sides of the profile.

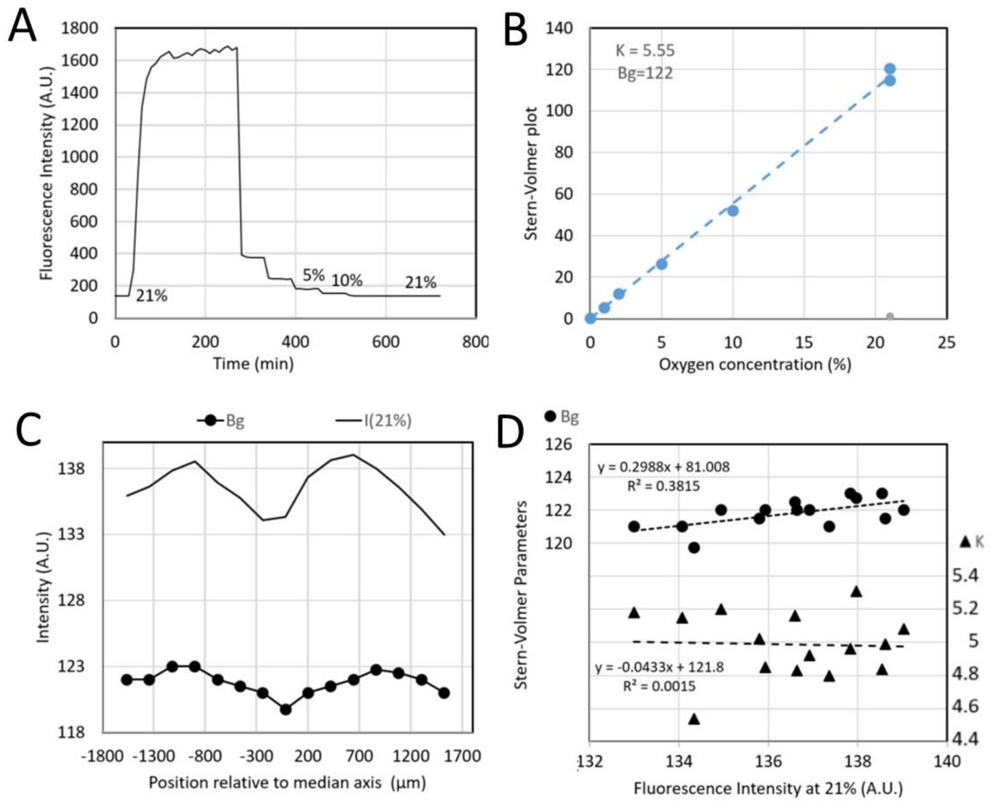


**Fig. S7.** Typical calibration data of sensing films mounted on a microfluidic device. A) Fluorescence intensity changes when applying an oxygen concentration ramp with the concentration in each gas channel. B) Corresponding Stern-Volmer plot (see text). C) Measured intensity at 21% $O_2$ level (solid line) and fitted background $B_g$ (bullets) for different ROI locations depicted in yellow in Fig. S6B. D) Fitted $B_g$ (bullets) and Stern-Volmer sensitivity parameter $K$ (triangles) as a function of 21% $O_2$ level intensity.

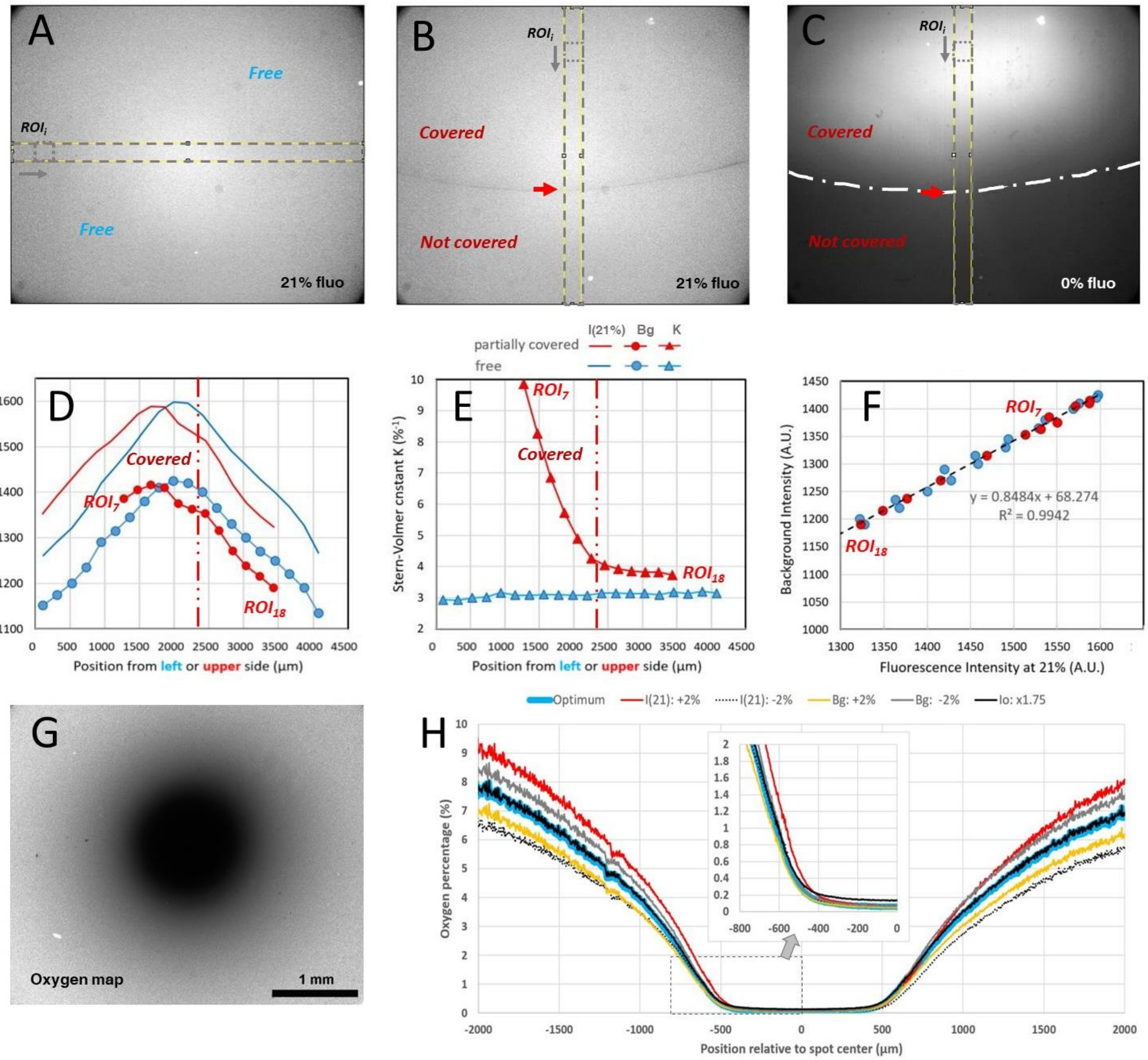


**Fig. S8.** Typical calibration data and oxygen profile measurement with covered sensing films for the spot assay. A) Homogeneous 21% $O_2$ level fluorescence image of an uncovered sensing film. B) Homogeneous 21% $O_2$ level fluorescence image of sensing film partially covered by a plain coverglass simulating the border of the spot assay (the red arrow indicates the glass boundary also depicted as a long dotted line in C). C) Fluorescence intensity when pure $N_2$ was flushed for 80 min (same region as in B) with the partial coverage of the sensing film). D) Measured intensity at 21% $O_2$ level (solid line) and fitted background $B_g$ (bullets) for different ROI positions depicted in yellow in A) for uncovered film (blue color) and in B) for partially covered film (red color). E) Fitted $B_g$ (bullets) and Stern-Volmer sensitivity parameter $K$ (triangle) along the ROI positions for the uncovered (blue color) and partially covered situation (red color). F) Fitted $B_g$ plotted as a function of the 21% $O_2$ level intensity for the uncovered and covered situations. G-H) Calculated $O_2$ map of *a Dictyostelium* spot covered and corresponding profile along a median horizontal line. Colors correspond to slight changes within the experimental uncertainty of the parameters used for the calculation (see text), insets correspond to the hypoxic (~0-2%) left side of the profile.

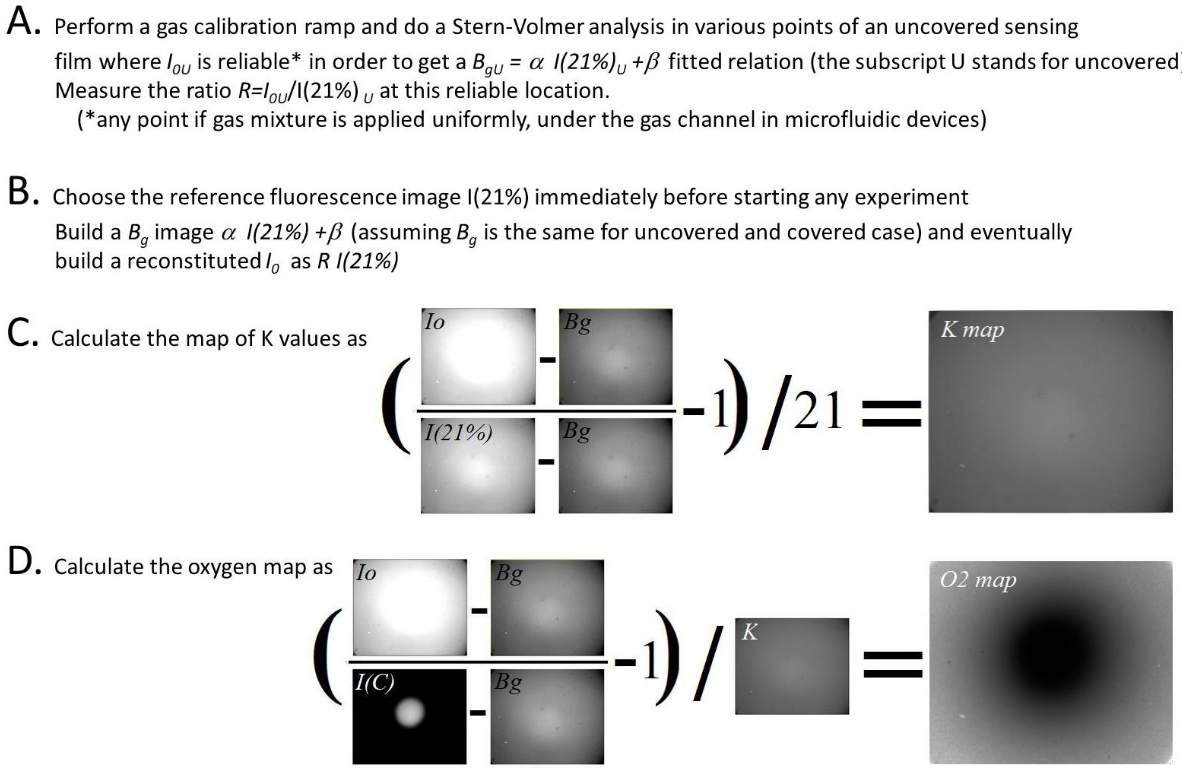


**Fig. S9.** Image analysis pipeline to quantify oxygen map from $O_2$ sensitive sensing films. Images correspond to an hypoxic circular zone created by a confined *Dictyostelium* spot.

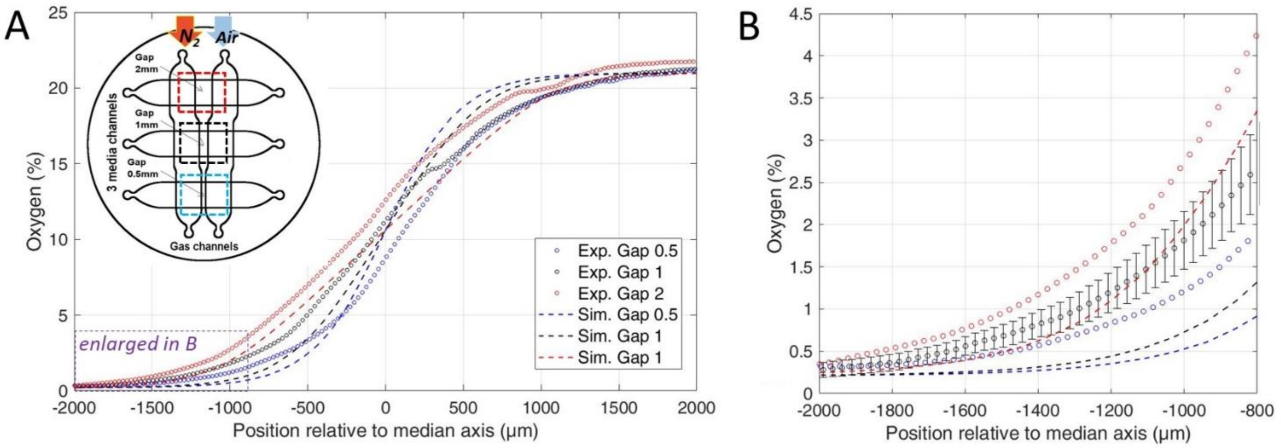


**Fig S10**. A) Comparison of the measured stationary oxygen profile in the microfluidic device (circles) and simulated ones (dotted lines) for the three gaps. Oxygen is measured thanks to the sensing film. The inset shows the gas injection conditions in the device: pure $N_2$ and air are flushed in left and right gas channel respectively. B) Enlarged oxygen profile in the hypoxic side. The estimated error bar on experimental measurements (showed for clarity on gap 1 mm data only) is explained in section 1.8 of the supporting information. Simulations are made with Comsol and explained in section 2.1 of the supporting information.

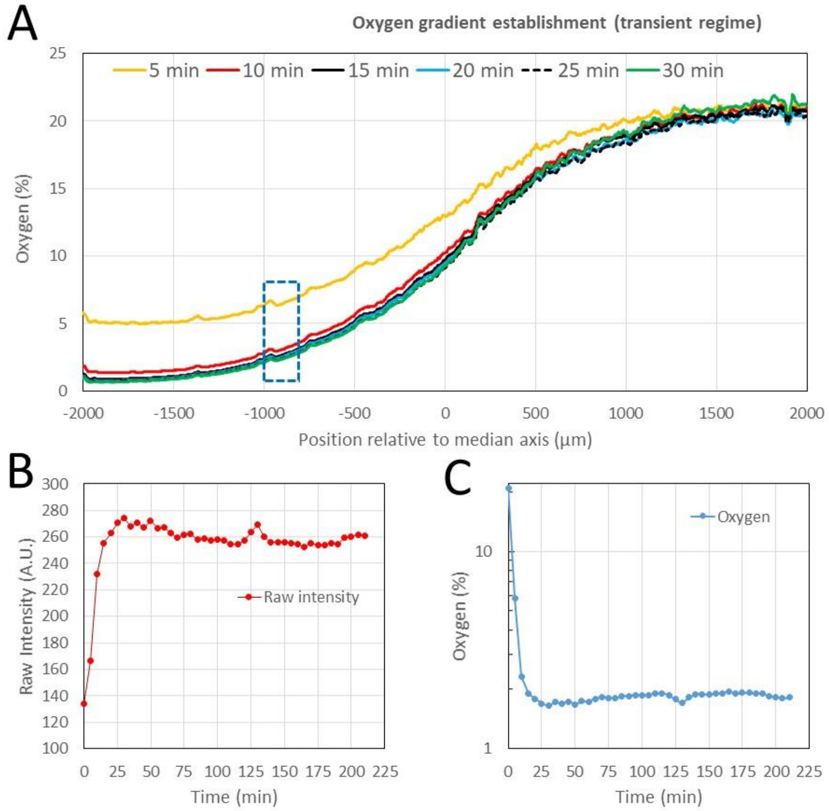


**Fig. S11.** Experimental oxygen gradient establishment in the microfluidic device (gap 0.5 mm). Pure $N_2$ and air are flushed at time 0 in left and right gas channel respectively. A) Oxygen profile measured using the sensing films at 5-min time interval. B-C) Raw intensity and measured oxygen in the ROI between -1000µm and -800µm from the device median axis in the region where the oxygen is about 1.5% (dotted region in A). Within 15 min, each signal reached 95% of its equilibrium value.

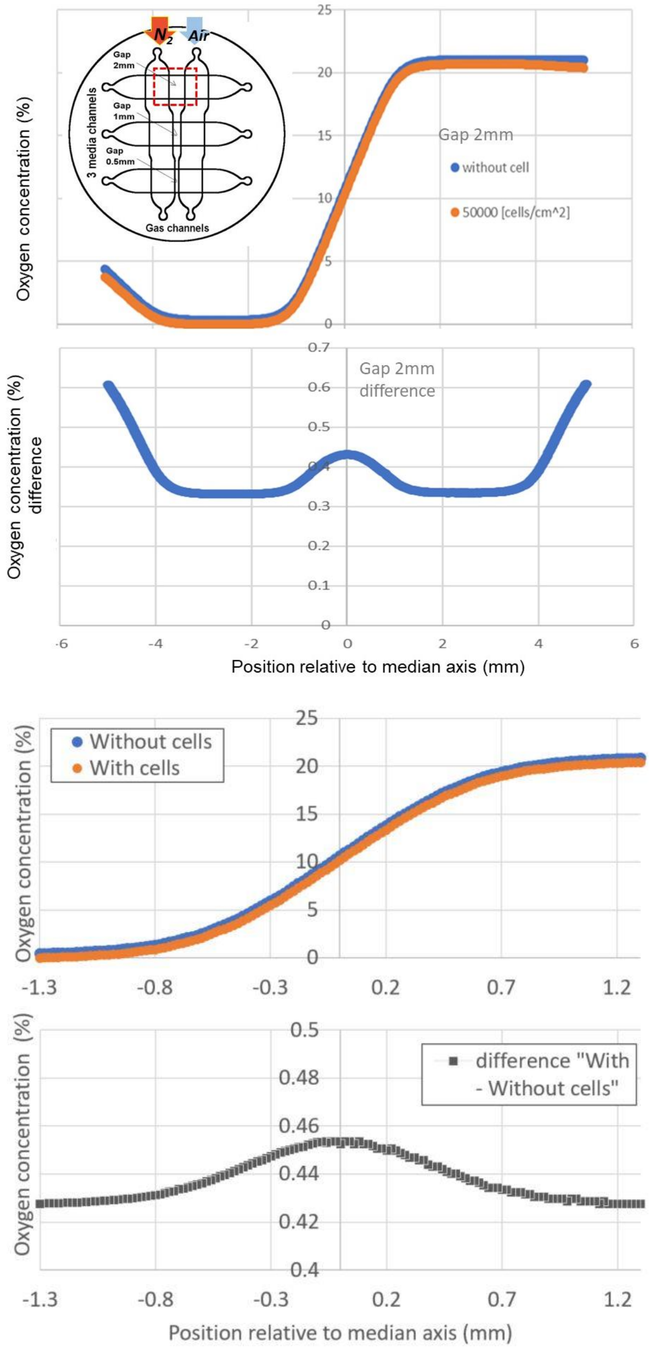


**Fig. S12. Influence of plated cells on the steady oxygen tension in the microfluidic device (Computational results)**. A) Absolute value of the oxygen concentration as a function of the position relative to the median axis of the device for the gap 1mm channel in presence (orange markers) or absence of cells (blue markers). Nitrogen is supplied on the left gas channel and 21% $O_2$ is supplied on the right channel. Cells density was taken as 500 cells/mm$^2$, which is the upper experimental limit. B) Corresponding difference between the two simulated situations (presence and absence of cells). In the region of interest where cells exhibit a strong aerotactic response (*i.e.*, around 1% $O_2$ or -1mm from the median axis), this difference is around 0.43% $O_2$ which is comparable to the error bar on $O_2$ measurements using sensing films (Fig. 2B). The rate of oxygen consumption by *Dd* cells was taken as 1.2.10$^{-16}$ mol.cell$^{-1}$.s$^{-1}$. Oxygen transport or consumption values or device dimensions are given in supplementary Table ST1 and in supporting information sections 1.4 and 2.1 respectively.

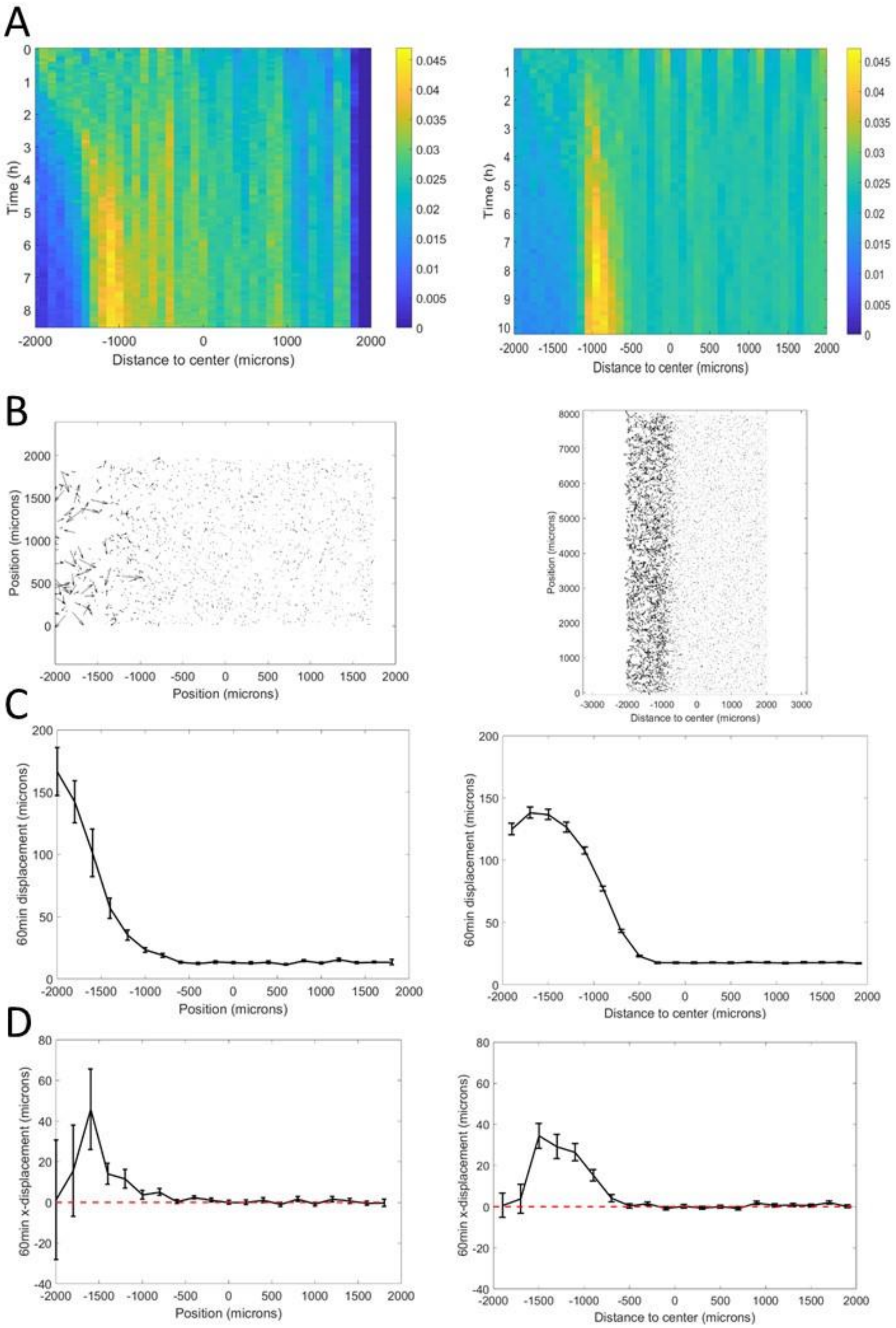


**Fig S13**. Adjusting Potts model (right) to microfluidic experiments (left). A: density kemographs showing cell accumulation in a low oxygen region. Colorbar represents the fraction of all cells within a bin in distance. B: Vectorial displacements displayed by cells over 1h. Higher activity at low oxygen concentrations, on the left, is clearly visible in both cases. C: quantification of cell activity as the norm of the displacements over 1h at different positions in the central channel. D: quantification of cell bias as the displacement of the cells in the x-direction over 1h.

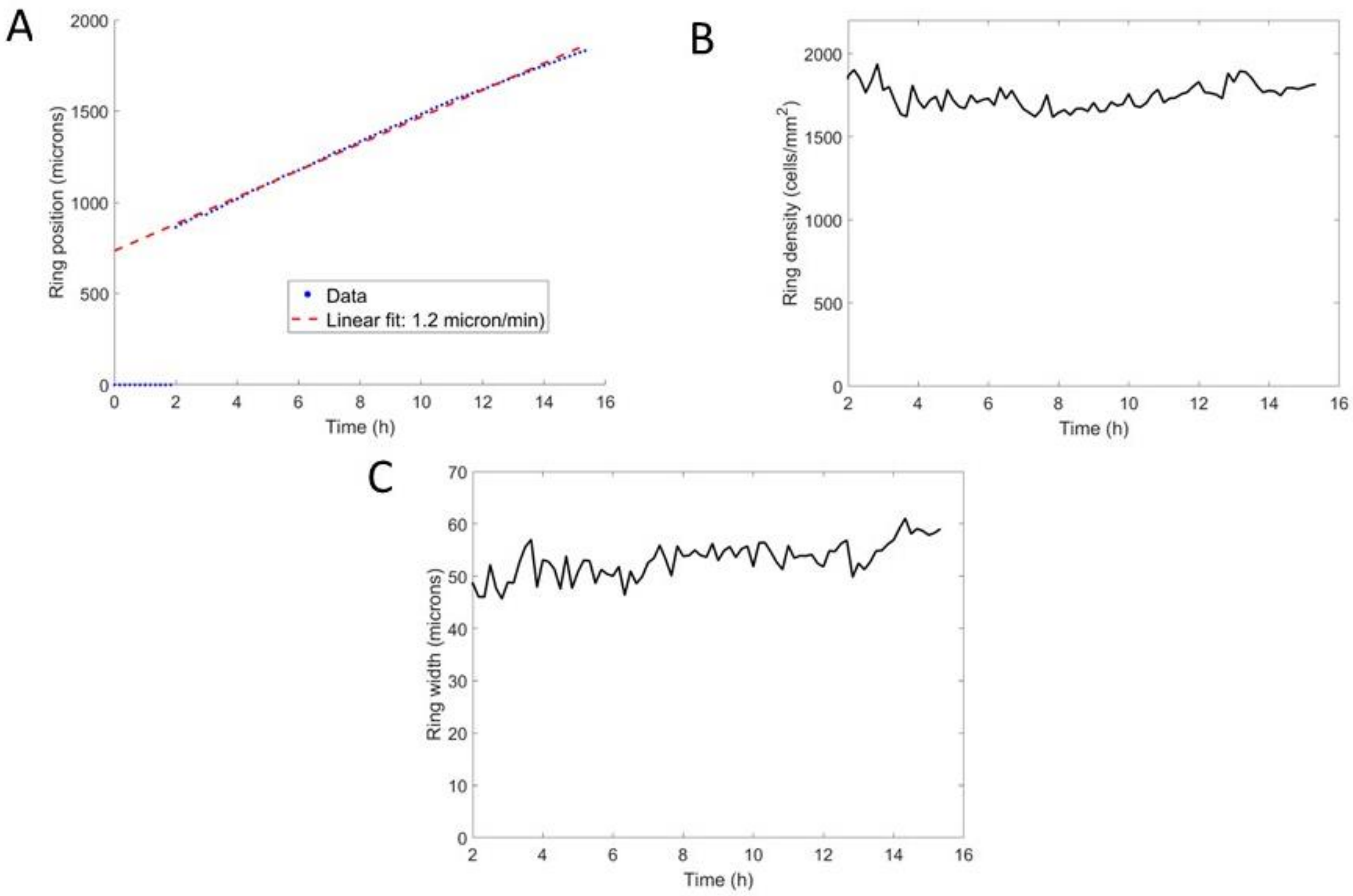


**Fig S14**. Potts model ring features with parameters adjusted from the microfluidic experiments (Fig S13). A: position of the ring as a function of time along with a linear fit yielding a speed of 1.1 µm/min. B: cell density within the ring as a function of time, after its formation. C: width of the ring as a function of time.

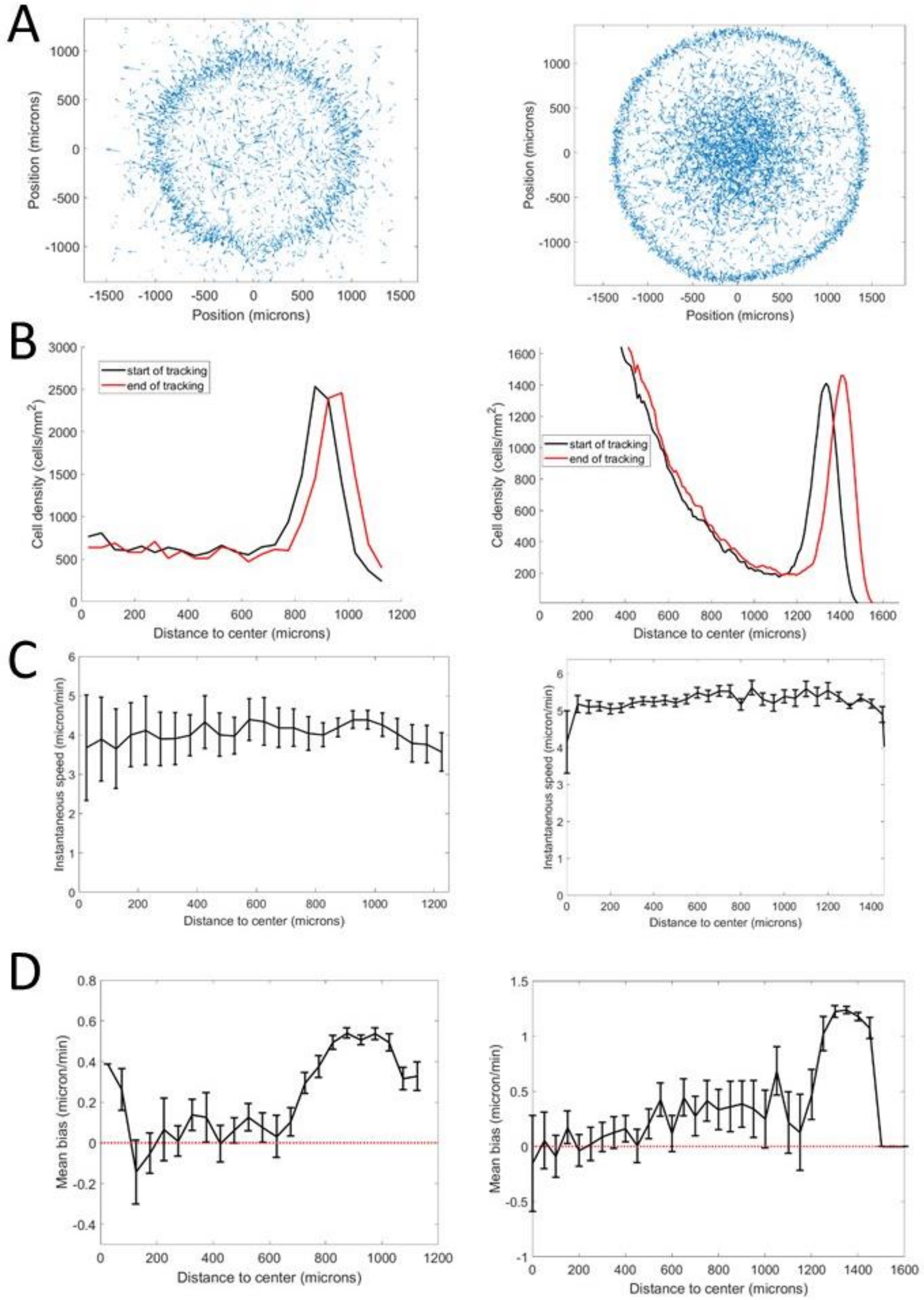


**Fig S15.** Comparison of cell behavior in spot experiments (left) and Potts models (right). A: Sample vectorial displacements over 1h. B: Cell density profiles at the beginning (black) and end (red) of the time window used for cell tracking. This indicates ring position for other plots. C: quantification of cell velocities as a function of position. D: Mean bias in the radial direction measured as the norm of the projected velocity in that direction. All error bars represent mean±std in each bin.

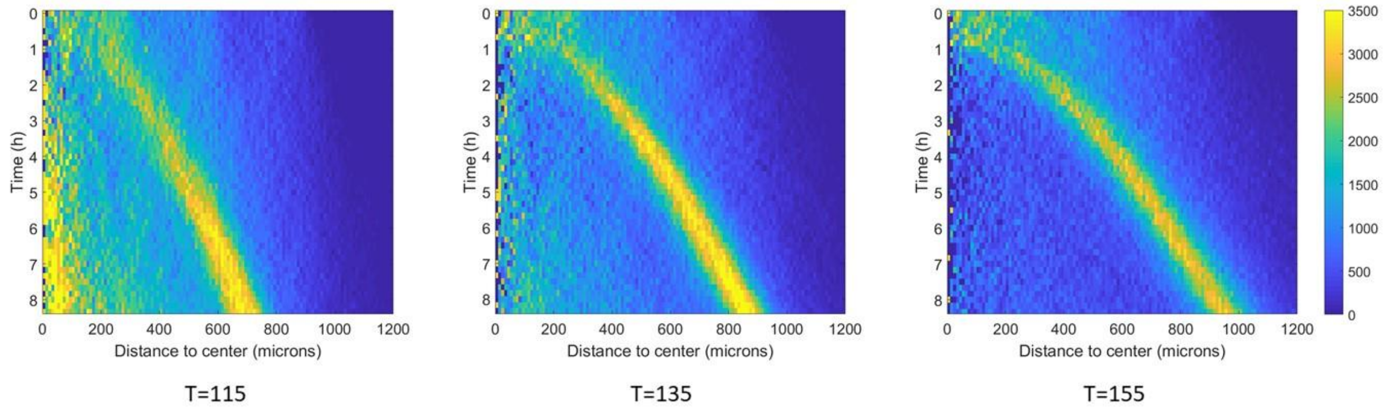


**Fig S16.** Effect of temperature on ring migration in Potts models. Density kemographs (all similarly scaled) of the full model, deprived of aerokinesis, at three different temperatures. Colorbar represents cell density in cells/mm$^2$. As temperature increases, fewer cells are left behind the ring and the ring moves at faster velocities.

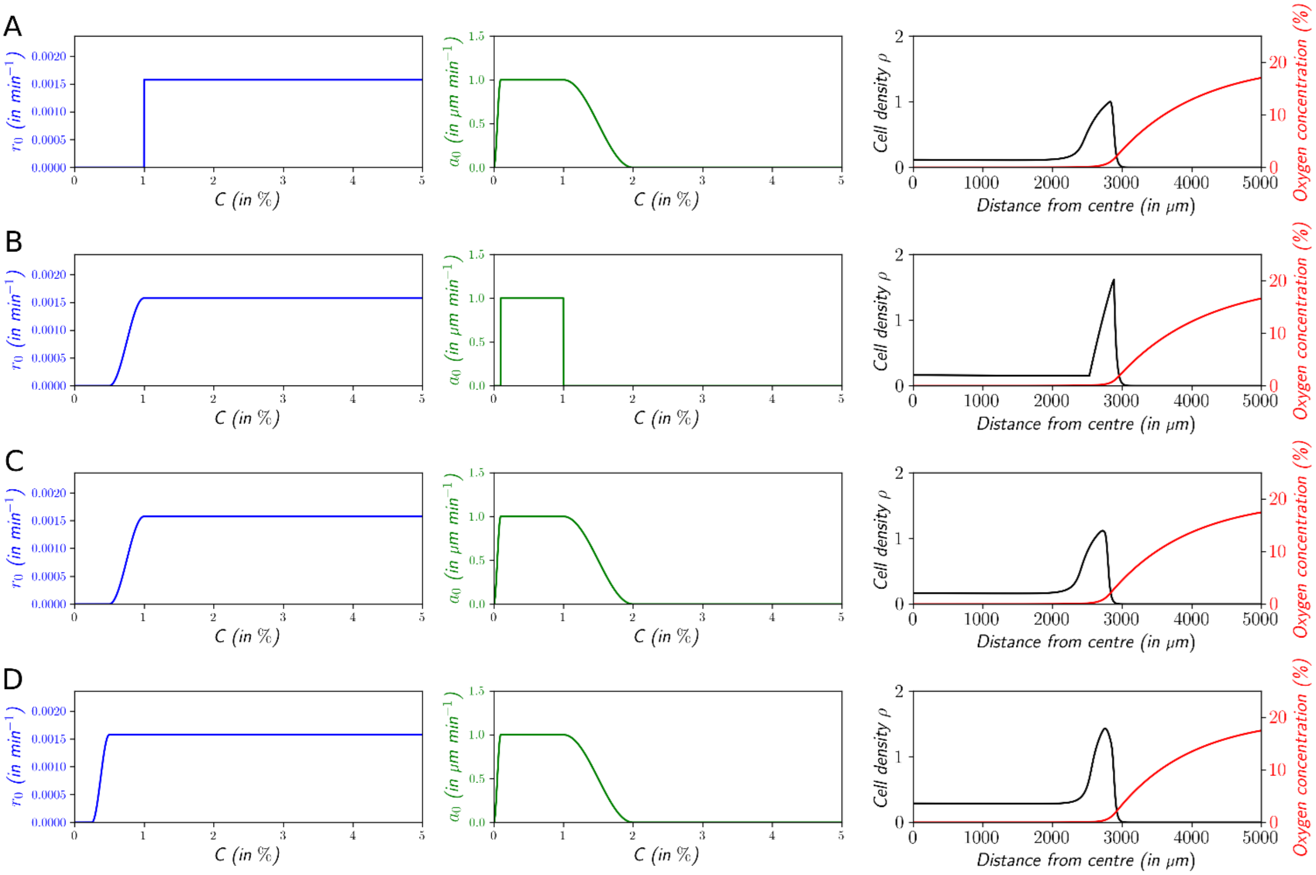


**Fig S17.** The shape of the profile and the speed of the wave vary according to the precise shape of the advection speed function $a(C)$ and the cell division rate function $r(C)$. A : $\sigma = 1.09 \mu m \cdot min^{-1}$, B : $\sigma = 1.03 \mu m \cdot min^{-1}$, C : $\sigma = 1.13 \mu m \cdot min^{-1}$, D : $\sigma = 1.19 \mu m \cdot min^{-1}$

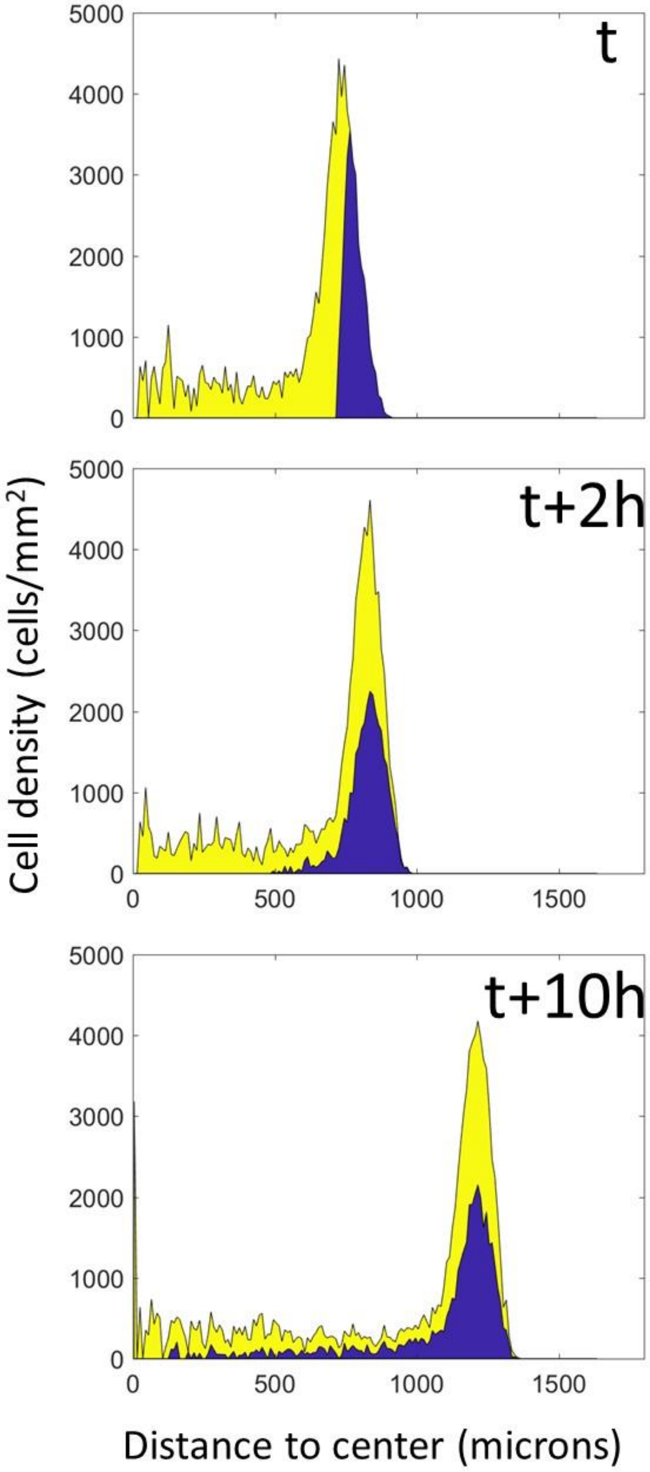


**Fig S18.** Mixing in Potts models. A simulation of the full model was stopped at an arbitrary time and cells were colored according to their position at this time. The simulation was then restarted from this time point and left to run for the equivalent of 10h, at which point complete mixing of both populations is clearly visible.

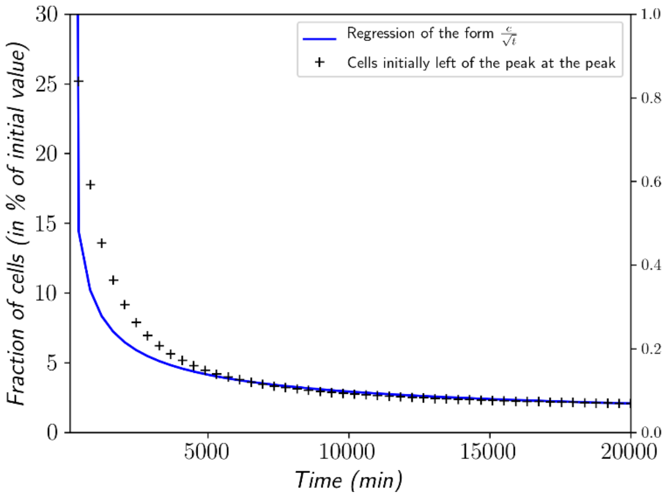


**Fig S19**. Decay rate of the neutral fraction at the peak composed of all the cells initially left of the peak in the go or grow model.

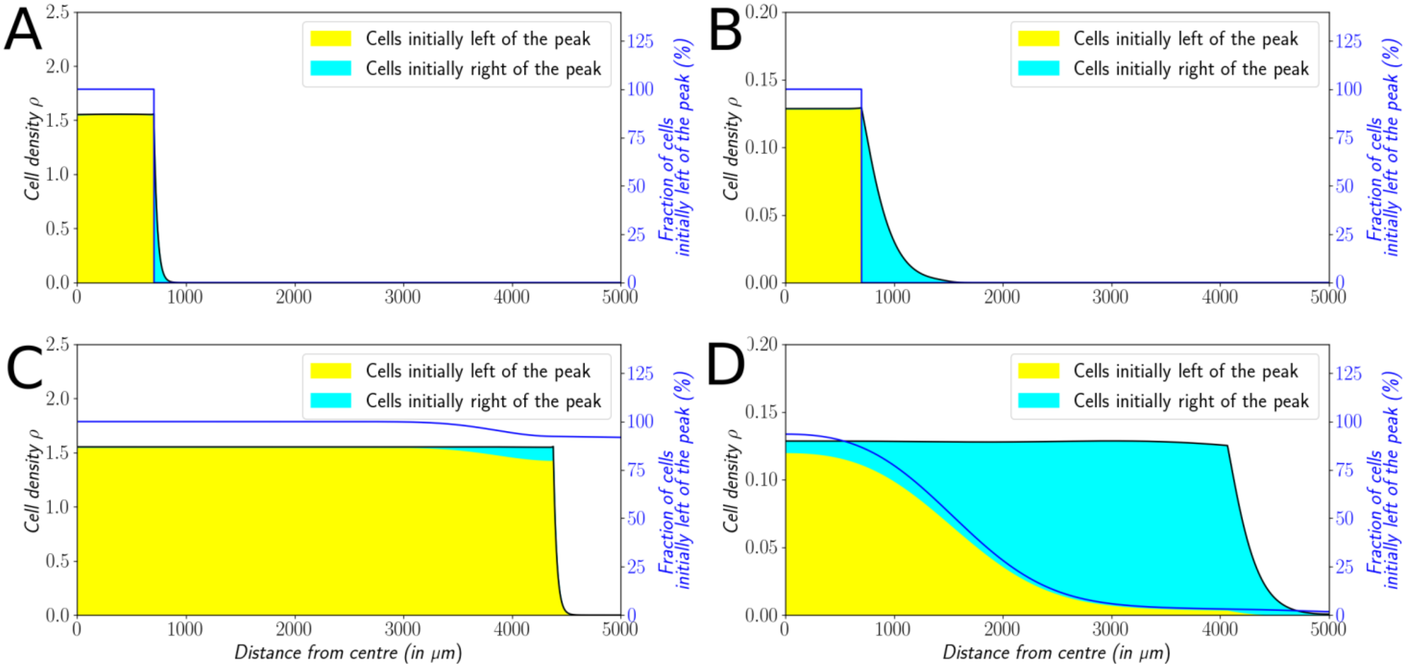


**Fig S20.** Cells initially left or right of the peak get labeled differently (A&B) in the single-threshold model. The labeling is neutral and does not change the dynamics of the cells. We let evolve the two colored population for some time and observe the mixing of the colors (C&D). A&C : With $a_0 = 1\mu m \cdot min^{-1}$, the wave is a pushed wave and after some time the front undergoes a spatially uniform mixing. B&D : With $a_0 = 0.1\mu m \cdot min^{-1}$, the wave is pulled and only the fraction initially in the front is conserved in the front.

#### 5. Captions for Supplementary movies

Movie M1. Initial phase (0-4h) of ring formation and migration. Scale bar: 500μm.

Movie M2. High framerate, high resolution imaging of cell dynamics in and behind the ring over 15min. Time is in minutes:seconds and the scale bar represents 100μm.

Movie M3. Reconstruction of cell and oxygen dynamics from a spot experiment on an oxygen sensor. Cell positions are shown as black dots, oxygen in colors (scale bar in %). The entire movie spans 15h of experiment.

Movie M4. Dynamics of the Potts model reproducing microfluidic experiments. Low oxygen regions are on the left and high oxygen on the right. Cell positions are shown as black dots and the entire movie represents the equivalent of 10h of experiments.

Movie M5. Dynamics of the Potts model reproducing the spot experiments. Cell positions are shown as black dots and the oxygen is in colors (in %).
